# Cholestasis alters polarization and suppressor function of hepatic regulatory T cells

**DOI:** 10.1101/2024.05.17.594680

**Authors:** Ramesh Kudira, Zi F Yang, Immaculeta Osuji, Rebekah Karns, Priya Bariya, Liva Pfuhler, Mary Mullen, Amy Taylor, Hong Ji, Celine S Lages, Annika Yang vom Hofe, Tiffany Shi, Srikar Pasula, Joseph A Wayman, Anas Bernieh, Wujuan Zhang, Claire A Chougnet, David A Hildeman, Gregory M Tiao, Stacey S Huppert, Sanjay Subramanian, Nathan Salomonis, Emily R Miraldi, Alexander G Miethke

## Abstract

Fibrosing cholangiopathies, including biliary atresia and primary sclerosing cholangitis, involve immune-mediated bile duct epithelial injury and hepatic bile acid (BA) retention (cholestasis). Regulatory T-cells (Tregs) can prevent auto-reactive lymphocyte activation, yet the effects of BA on this CD4 lymphocyte subset are unknown. Gene regulatory networks for hepatic CD4 lymphocytes in a murine cholestasis model revealed Tregs are polarized to Th17 during cholestasis. Following bile duct ligation, *Stat3* deletion in CD4 lymphocytes preserved hepatic Treg responses. While pharmacological reduction of hepatic BA in MDR2^-/*-*^ mice prompted Treg expansion and diminished liver injury, this improvement subsided with Treg depletion. A cluster of patients diagnosed with biliary atresia showed both increased hepatic Treg responses and improved 2-year native liver survival, supporting that Tregs might protect against neonatal bile duct obstruction. Together, these findings suggest liver BA determine Treg function and should be considered as a therapeutic target to restore protective hepatic immune responses.

## Introduction

Fibrosing cholangiopathies in children and young adults include extrahepatic biliary atresia (EHBA) and primary sclerosing cholangitis (PSC). They result from immune-mediated injury to the intra- and extrahepatic biliary tree, sterile inflammation, and biliary fibrosis which progresses rapidly to cirrhosis in infants with EHBA. In many patients, complications of cirrhosis necessitate liver transplantation to prolong survival ^1–4^. Studies of the genome and liver transcriptome in patients diagnosed with a cholangiopathy revealed a regulatory connectome involving the genes *STAT3, IL6, TNFa* and *FOXP3* controlling cellular immunity and fibrosis ^5^.

Investigations on liver and bile duct tissue from patients with EHBA or PSC point at an imbalance between Tregs and effector lymphocytes, including cytotoxic CD8 and IL17A producing CD4 T-cells, as drivers in the pathogenesis of both diseases ^6–10^. Epithelial infiltration of intrahepatic bile ducts and oligoclonal expansion of CD8 T-cells isolated from biliary remnants demonstrate a key role of CD8 lymphocytes in initiating EHBA ^11, 12^. Similarly, a high prevalence of liver infiltrating CD28^neg^ CD8 lymphocytes that escape control by co-stimulation and a dominant network of CD8 T-cells and neutrophils isolated from the biliary epithelium suggest effector functions for CD8 lymphocytes in the pathogenesis of PSC ^8, 13^. In animal models for both diseases, including the MDR2^-/-^ mouse recapitulating small duct PSC and rhesus rotavirus (RRV)-induced experimental EHBA, targeting CD8 lymphocytes with depleting antibodies attenuated the disease phenotype ^6, 10, 14, 15^. Produced by lymphocytes, IL17A has been recognized as a chief cytokine in directing immune mediated injury to the biliary epithelium in both conditions ^7, 16^.

Tregs are a small subset of CD4 lymphocytes expressing the master transcription factor FOXP3 and the surface receptor IL2RA (CD25) ^17^. They are implicated in controlling bile duct obstruction in experimental EHBA ^18–20^. A polymorphism in the *IL2RA* gene is associated with decreased number and impaired function of Tregs in patients with PSC ^21^. Tregs are indispensable for maintaining peripheral tolerance and preventing autoimmunity in antigen-dependent fashion ^22^. The role of Tregs in restoring tissue integrity and repair following epithelial injury are increasingly recognized ^23^. These innate functions are mediated, at least in part, by secretion of amphiregulin (AREG), the ligand of epidermal growth factor receptor (EGFR). Immunotherapies boosting Tregs to prevent autoimmunity and to promote epithelial repair and regeneration are at various clinical trial stages. Preliminary results from phase 1 and 2 studies demonstrate the efficacy and safety of low-dose interleukin (IL)-2 treatment for Treg expansion in SLE and IBD ^24–26^. Our group has shown that treatment with IL-2 immune complexes promotes hepatic Treg expansion reduces the number of CD8 lymphocytes, and halts biliary injury progression in MDR2^-/-^ mice ^6^.

Under physiological conditions, primary bile acids (BA) like cholic or chenodeoxycholic acids (CA or CDCA) are synthetized from cholesterol in hepatocytes, conjugated with taurine or glycine, and transported across the canalicular membrane by BSEP. They are then excreted with bile into the intestine, where they facilitate absorption of fats and fat-soluble vitamins. 95% of bile acids are reabsorbed in the terminal ileum. The transportation of BA is facilitated by ileal bile acid transporter (IBAT). The inhibition of IBAT using small molecules has been shown to result in anti-cholestatic effects ^27^. Upon injury to the biliary tree in PSC or EHBA, excretion of conjugated BA like TCDCA, TCA, or GCDCA into bile is diminished, leading to their retention in liver and plasma ^28^. Defective biliary excretion of phospholipids due to deletion of *Abcb4* (MDR2) results in the precipitation of microcrystals in bile canaliculi, injuring hepatocytes and interlobular bile ducts. This causes biliary fibrosis and highly elevated serum and liver BA concentrations, especially in BALB/c mice ^29–31^. Regurgitation of toxic BA from leaky bile ducts into the periductal space produces sterile inflammation in these mice ^14^. In contrast to this model of chronic cholestasis and sclerosing cholangitis (SC), treatment with xenobiotic 3,5-diethoxycarbonyl-1,4-dihydrocollidine (DDC) and bile duct ligation (BDL) are models of acute SC, with BDL representing the purest form of BA-induced hepatobiliary injury ^32–35^.

Primary BA were recently described to modulate cytokine production by monocytes in children with chronic cholestasis, but their effects on hepatic T lymphocytes remain largely undefined ^36^. EHBA is the cholangiopathy with the highest reported serum BA levels, of which TCA and TCDCA are the most abundant conjugates and are increased by 40- and 23-fold, respectively, compared with age-matched healthy controls exceeding serum BA concentrations of 100 μM at diagnosis ^37^. Unless bile flow in EHBA is surgically restored by hepatic portoenterostomy (HPE), liver transplantation within the first 2 years of life is inevitable. If the procedure is performed within the first 90 days of life, 2-year survival with native liver (2yrSNL), an established clinical endpoint, is achieved in approximately 53% of infants ^38^. Here, we examine the effects of BA on Treg homeostasis in preclinical models of acute and chronic SC and in infants with EHBA.

## Results

### Hydrophobic bile acids repress FOXP3 and regulatory T cell function

In vitro studies were performed to determine the effects of primary BA on Treg molecular phenotype and function. Hydrophobic CDCA, but not hydrophilic CA, repressed FOXP3 in cultured splenic Tregs (Fig 1A). When used at concentrations found in the plasma of infants with EHBA^37^, taurine conjugates of CDCA reduced FOXP3 expression in purified Tregs from adult and neonatal mice (Fig 1B/C) and repressed *Foxp3* transcription (Fig S1A) without significantly compromising the viability or inducing apoptosis in these CD4 cells (Fig S1A-D). CDCA and TCDCA also reduced in vitro Treg induction in hepatic and splenic naïve CD4 cells (Fig 1D, Fig S1E). Activation of Tregs in media conditioned with TCDCA decreased the expression of Treg suppressor molecules, including CD39, CTLA4, PD-L1, and AREG (Fig 1E, Fig S1G) and diminished their capacity to inhibit proliferation of CD8 lymphocytes in suppressor co-cultures assays (Fig 1F, Fig S1H). TCDCA repressed FoxP3 expression in cultured Tregs separated from buffy coat of healthy blood donors and CDCA diminished Treg-induction in naïve CD4 cells from these donors (Fig 1G, Fig S1F). To assess the effects of BA on hepatic Treg homeostasis in a model of acute cholestasis and sterile inflammation, mice were treated for 14 days with 0.1% (wt/wt) of DDC, which increased serum markers of hepatocellular (ALT) and duct injury (ALP) and biomarkers of cholestasis (total bilirubin [TB], and liver/serum BA; Fig 1H/I). When DDC chow was restored to regular chow, cholestasis and hepatobiliary injury promptly resolved, evidenced by a rapid decline in both liver biochemistries and serum BA levels between D14+7 and D14+28 (Fig 1H/I). Rise in serum markers of liver injury during DDC challenge was accompanied by progressive SC on review of liver sections stained with hematoxylin and eosin (H&E), Sirius red (SR), and by immunohistochemistry (IHC) for CK19, involving periportal inflammation, biliary fibrosis, and ductal proliferation, respectively (Fig S2A). DDC-induced cholestatic liver injury was associated with a progressive decline in frequency of hepatic Tregs, which recovered in number when DDC challenge was discontinued (Fig 1J, Fig S2B). CD8+ lymphocytes infiltrated the liver upon DDC challenge and contracted afterwards (Fig S2B). In DDC-treated mice following Treg depletion via αCD25-antibody, the number of liver infiltrating CD8 cells increased, serum levels of ALT, ALP, and TB were elevated, and inflammation and ductal proliferation accelerated compared with isotype-IgG treated controls (Fig 1K/L, Fig S2C-F), suggesting hepatic Tregs play a protective role. In summary, conjugated hydrophobic BA repressed FOXP3 expression in Tregs and their suppressor capacity in vitro. In vivo, severe cholestasis from DDC-induced hepatobiliary injury was accompanied by a contraction of hepatic Tregs, a CD4 cell population which conferred protection from liver injury.

**Figure 1.**
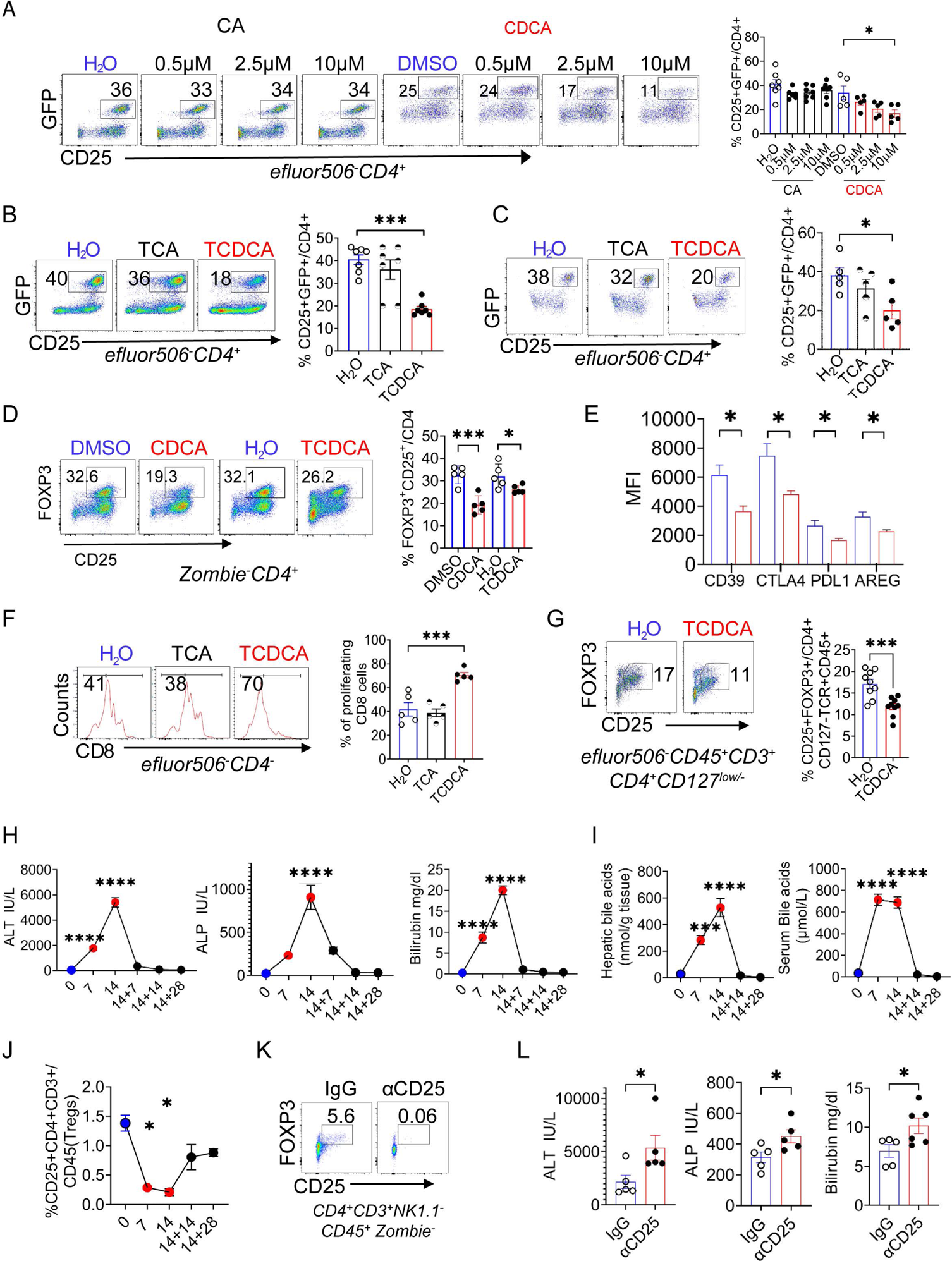
TCDCA represses FOXP3 in Tregs and reduces their suppressor function. Splenic Tregs from Foxp3/GFP reporter mice were stimulated with aCD3/CD28 and cultured with IL-2 media conditioned with various BA for 48 hours prior to cytometric enumeration of CD25+GFP (FOXP3)+ Tregs (**A**/**B**). Splenic Tregs from neonatal mice were cultured with 100µM of TCDCA or 100µM of TCA (**C**). Hepatic naïve CD62L positive CD4 cells were cultured with TGFb1, IL-2, aCD3/CD28 and stimulated with 10µM of CDCA or 100µM of TCDCA for 3 days (**D**). Tregs were stimulated with 100 µM of TCDCA and characterized via flow cytometry (**E**). Suppressor function of Tregs treated with 100µM TCDCA/TCA was assessed in proliferation assays with CD8 lymphocytes (**F**). FACS purified Tregs from healthy blood donors were treated with 100µM TCDCA (**G**). Mice were fed 0.1% DDC chow to induce SC; serum and tissue samples were collected at D0, during DDC challenge (D7/D14), and after (D14+14/28) to measure serum biochemistries (**H**), serum and liver BA concentrations (**I**), and to enumerate hepatic Tregs by flow cytometry **(J).** Tregs were depleted with αCD25 antibodies prior to a 7-day DDC challenge; hepatic Treg-frequency was measured with flow cytometry **(K)** and serum biochemistries were measured with colorimetric assays **(L)**. In **A-G**, the data points represent results of individual wells from at least 2 independent experiments. In **H-J**, mean and SEM are represented for at least 4 mice/time point and in **L**, the data points represent results from individual mice. Multiplicity adjusted *p*-values were computed using one-way ANOVA with Dunnett’s post-hoc test compared to mean of the vehicle in **A, B, C, F** or the mean of day 0 baseline values in **H-I** with \**p*<0.05; \*\**p*<0.01; and \*\*\**p*<0.001. Unpaired t-tests were applied to determine significant differences in **D, G** and **L**. See also Figures S1 and S2.

### Transcriptional profiling of hepatic Tregs with single cell genomics predicts cholestasis induced Th17 polarization

To better understand transcriptional regulation of hepatic Tregs under cholestatic conditions, we conducted single-cell (sc)RNA-seq and scATAC-seq to characterize the gene expression and chromatin accessibility changes, respectively, in parenchymal and non-parenchymal liver cells from mice before, during, and after DDC challenge (Fig S3A). scRNA-seq resulted in 101,044 quality cells, from which we identified 16 cell populations, spanning epithelial, endothelial, and immune cells (Fig 2A). Sub-clustering of T cells and innate lymphoid cells (ILCs) yielded a population of NK cells, Type 1 ILCs (ILC1) and 8 distinct T cell populations: naïve T cells (*Ccr7, Il7r, Tcf7, Sell*), effector CD8 T cells (*Cd8a, Gzma*), two CD4 tissue-resident memory (TRM) T cell populations (*Cd4, Cd69, Cxcr6, Cxcr3*), Tregs (*Cd4, Ctla4, Foxp3*), interferon-stimulated T cells (*Ifng, Isg15, Isg20, Ifit1, Ifi3*), and gamma-delta T(gdT) cells (*Trdc*) (Fig 2B, D, Fig S3B). Among the regulatory and effector memory CD4+ populations, we identified 879 upregulated genes that showed cell-type and/or time-dependence (Fig 2E, FDR = 10%, log2(FC) > 1, Tables S1 and S2). *Foxp3* and *Ctla4* were uniquely upregulated in Tregs relative to the other CD4+ T cell populations. Furthermore, in response to DDC injury, repair gene *Areg* and Th17 signature genes *Il23r*, *Il17a*, and *Rorc* were uniquely and dynamically induced in Treg (Fig. S3D). Surveying the transcriptome of Tregs for canonical transporters of conjugated BA, we detected no expression for NTCP (*Slc10a1*) or ASBT (*Slc10a2*), or their surface and intracellular receptors, TGR5 (*Gpbar1*) and FXR (*Nr1h4*), respectively. However, sphingosine-1-phosphate receptors (*S1pr1, S1pr4*) were enriched in Tregs (Fig 2G). Of note, conjugated BA were reported to activate S1PR2 on rat hepatocytes and cholangiocarcinoma cells, raising the possibility that S1PRs mediate effects of TCDCA on Tregs^39, 40^. Of putative Treg suppressor molecules, *Ctla4 and Pdcd1* were most abundantly expressed by hepatic Tregs in the context of DDC-induced SC (Fig 2H). To explore underlying gene regulatory mechanisms, we performed scATAC-seq. Our analysis identified 20 cell populations from 41,387 quality cells, including 7 T cell subsets (Fig 2C, Fig S3C). Of 330,504 total accessible regions (peaks), 312,061 were differentially accessible across populations (Fig 2F, FDR = 10%, log2|FC| > 1). Among the T lymphocyte populations, the *Foxp3* promoter and putative intronic enhancers regions were uniquely accessible in Tregs, while accessible regions in the *Areg* and *Ctla4* loci had elevated accessibility in Tregs, relative to the other CD4 and γοT cell populations.

**Figure 2.**
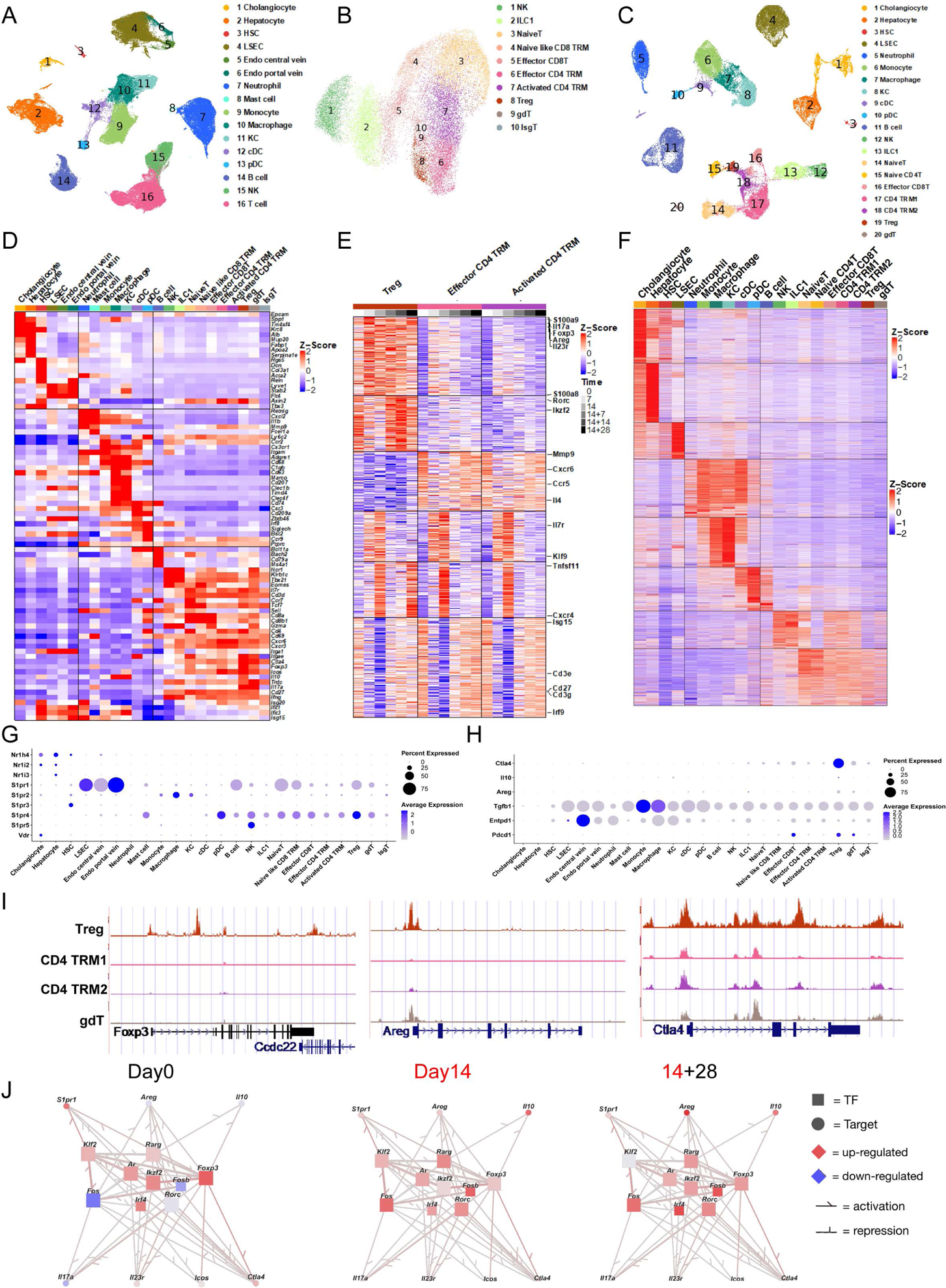
Gene regulatory networks in hepatic Tregs during progressive cholestasis. scRNA-seq was performed on single-cell suspensions from mononuclear cells and preparations of epithelial, endothelial, and hepatic stellate cells from livers of mice at D0 (n=3), D7(n=1), D14(n=2), D14+7(n=1), D14+14(n=3), D14+28(n=1). scRNA-seq revealed 16 major cell types displayed in the UMAP (**A**). Further analysis of NK cells and T lymphocytes identifies 10 cell subtypes (**B**). snATAC-seq was performed on single nuclei suspensions from livers of mice at D0(n=2), D14(n=1), D14+14(n=1), D14+28(n=1) of DDC challenge. snATACseq analysis found 20 cell types through scRNA-seq label transfer, shown in the peak based UMAP (**C**). Pseudobulk RNA heatmap displays cell type specific marker gene expression, z-scored across all cell types (**D**). Pseudobulk gene expression heatmap shows signature genes by cell types and time points in Treg and two CD4 T cell clusters with FDR cutoff 10%, Log2FC>1, z-scored across conditions. The top 10 highest expressed genes at each cell type and time point are annotated on the right (**E**). Pseudobulk ATAC-seq heatmap displays 312,061 differential peaks through DEseq2 comparison, FDR cutoff 10%, Log2|FC|>1 (**F**). Dot plot gene expression for putative bile acid receptors and Treg-associated suppressor molecules across all liver cell types from scRNA-seq (**G/H**). Peak tracks show locus accessibility for Foxp3 to be exclusive in Tregs (**I**). Accessibility for *Areg* is predominantly in Tregs, but all CD4 and gd T lymphocytes populations contain accessible chromatin for regulation of *Ctla4*. Gene regulatory networks in Treg demonstrate dynamic gene activation across time points (**J**). Transcription factors (TF) were colored by estimated TF activity, target genes were colored by gene expressions, z scored across Treg and two CD4 TRM cell clusters. See also Figure S3 and Table S1-3.

Combining scRNA-seq with scATAC-seq, we built gene regulatory networks (GRNs), to predict regulatory interactions between TFs and target genes in the CD4+ memory T cell populations of the liver (see Methods)^41, 42^. We predicted that Treg expression of Th17-associated genes (e.g., *Il17a*, *Il23r*) during DDC injury were regulated by Rorc, Fos and Fosb (Fig 2J, Table S3). Indeed, the expression of *Rorc*, *Fos*, and *Fosb* is correlated with their predicted Th17 targets (Fig. S3D). Members of the AP-1 complex, including c-Fos, Fosb, and Fosl2, were previously reported to control Th17 polarization, Il23p19 production, and IL17A induced CCL20 expression ^43–45^. In summary, we identified a population of hepatic Tregs in both the scRNA-seq and scATAC-seq datasets. Treg responded to cholestatic liver injury and tissue repair by upregulating Th17-associated cytokines and *Areg* under the control of TFs Rorc, Fos, and Fosb.

### Hypermethylation of the *Foxp3* promoter region mediates bile acid induced silencing of Treg responses

Following leads from the sc-genomic analyses, splenic Tregs from *Nr1h4* (FXR) and from *Gpbar1* (TGR5) deficient mice were cultured with conjugated BA. These cells were similarly susceptible to TCDCA-induced repression of FOXP3 as Tregs from WT mice, confirming that canonical BA receptors do not mediate the effects of TCDCA (Fig 3A, Fig S4A). In contrast, blocking of S1PR1 or S1PR4 with small molecule antagonists FTY720 or CYM50358, respectively, attenuated repression of FOXP3 in Tregs cultured with TCDCA in a dose-dependent fashion (Fig 3B/C, Fig S4B-D). We further showed that TCDCA induced RORγt expression in Tregs, which was repressed by blocking S1PR1 signaling (Fig 3D). It is well established that the methylation status of regulatory elements of *Foxp3,* which is critical for Treg-differentiation, is tightly controlled by the enzymes DNMT1 and DNMT3a ^46^. In culture, TCDCA induced mRNA expression of *Dnmt1* and *Dnmt3a* and increased relative protein concentrations of DNMT3a in splenic Tregs (Fig 3E/F). Pyrosequencing of DNA from cultured Tregs exposed to TCDCA revealed hypermethylated CpG islands in the *Foxp3* promoter region when compared with Tregs cultured in presence of vehicle or TCA (Fig 4G). Inhibition of DNMT activity with 5-Azacytidine (5-Aza) rescued Tregs from BA induced repression of FOXP3 (Fig 4H). When cholestatic MDR2^-/-^ mice were treated with 5-Aza, the proportion of Tregs among hepatic CD4 lymphocytes and the absolute number of liver infiltrating Tregs increased significantly (Fig 3I). This was accompanied by decreased hepatocellular injury with a trend to lower ALT levels, diminished periportal inflammation, and protection from biliary injury, as evidenced by lower serum ALP levels and reduced intrahepatic biliary mass by CK19 IHC (Fig 3J-L). 5-Aza treatment also increased hepatic CD8 cell number (Fig S4J). Using sex and age-matched MDR2^+/-^ littermates as non-cholestatic controls, 5-Aza treatment increased hepatic Treg number without discernable liver injury or liver histopathology (Fig 3I-L, Fig S4H/I).

**Figure 3.**
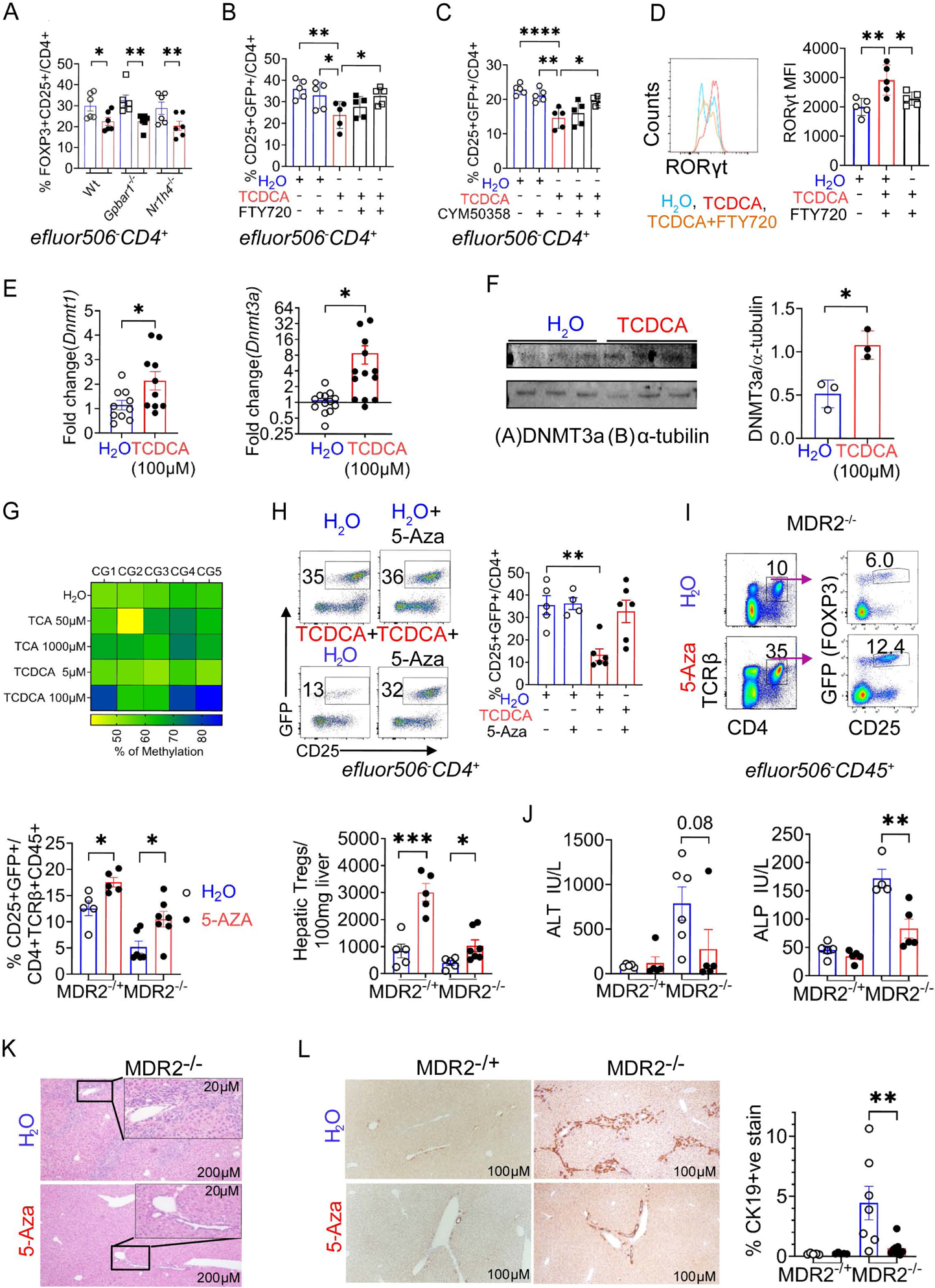
Effects of TCDCA on Tregs are mediated by S1P receptors and hypermethylation of the Foxp3 promoter region. Splenic Tregs from wild type (WT), *Gpbar1*(TGR5)-/-, and *Nr1h4*(FXR)-/- mice were cultured and stimulated with 100μM of TCDCA and percent FOXP3+ Tregs was enumerated by cytometry **(A).** Splenic Tregs from FOXP3/GFP reporter mice were cultured and stimulated with 100μM of TCDCA, 10/100 nM of the S1PR inhibitor FTY720, or 10nM/1μM of the S1PR4 inhibitor CYM50358 (**B/C**). Splenic Tregs were cultured and stimulated with 100μM of TCDCA ± 100nM of the S1PR1 inhibitor FTY720, followed by restimulation with PMA/Ionomycin and intracellular flow cytometry (**D**). Splenic Tregs were stimulated with 100μM of TCDCA for 24 hours prior to *qPCR* and immunoblotting for DNMT3a (95kDa) and a-tubulin (55kDa) with subsequent densitometry (**E/F**). Pyrosequencing analysis revealed percentage of methylation at five CpG sites of the *Foxp3* promoter DNA from cultured splenic Tregs stimulated with TCA or TCDCA. The heat map represents the mean values from two replicates of two independent experiments (**G**). Splenic Tregs were cultured and stimulated with 100μM of TCDCA and 1μM of the methylation blocker 5-aza-2’-deoxycytidine (5-Aza) or vehicle control for 5-Aza (H_2_O) (**H**). 20 ug/g mouse of 5-Aza or vehicle was administered by daily IP injections for seven days to cholestatic MDR2^-/-^ or MDR2^+/-^ mice serving as controls prior to cytometric enumeration of hepatic Tregs (**I)** Serum ALT and ALP levels at the end of 5-Aza treatment course (**J**). Representative photomicrographs of H&E-stained 5-Aza treated MDR2^-/-^ mouse liver sections (**K)**. CK19 immunohistochemistry and quantification from MDR2^-/-^ mice following 5-Aza treatment (**L**). Data points represent results from individual wells of at least two independent experiments in **A-E** and from individual mice in **I-L**. Differences between groups were tested for statistical difference by unpaired t-test in all panels except for **B and C** in which one-way ANOVA was applied with Dunnett’s multiple comparisons test against TCDCA without S1PR inhibitor as control with \**p*<0.05; \*\**p*<0.01; \*\*\*\**p*<0.0001. See also Figure S4

**Figure 4.**
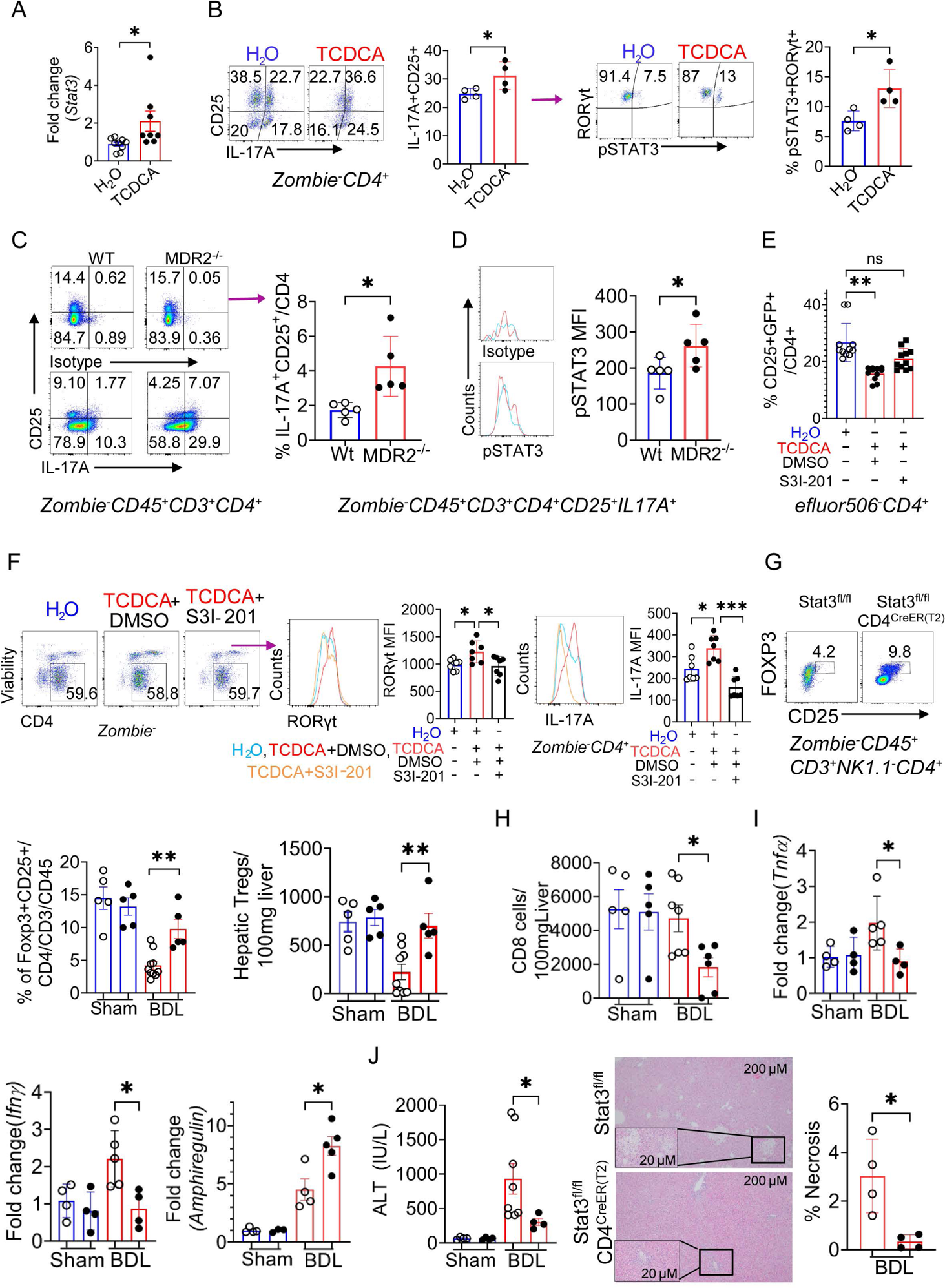
Deletion of *Stat3* in CD4 lymphocytes restores protective Treg-responses in cholestatic mice. Splenic Tregs were stimulated and cultured in presence of 100 μM of TCDCA or vehicle for 24 hours prior to qPCR or for 48 hours prior to restimulation with PMA/Ionomycin and intracellular flow cytometry (A/**B**). Liver mononuclear cells were isolated from WT or MDR2-/- mice and restimulated with PMA/Ionomycin prior to cytometry (**C**). FACS sorted splenic Tregs from Foxp3/GFP reporter mice were stimulated and cultured in presence of TCDCA or H_2_O in media conditioned with STAT3 antagonist S31-201(100μM) or DMSO (vehicle) (**D**). *Stat3*^fl/fl^ and *Stat3*^fl/fl^CD4^CreER(T2)^ mice were treated with tamoxifen prior to exposure to 0.1% DDC admixed to chow or control chow for 14 days prior to separation of liver mononuclear cells and cytometric enumeration of Tregs (**E**). MACS sorted splenic Tregs from WT mice were cultured in presence of 100µM of TCDCA or H_2_O in media conditioned with 100µM of S31-201 for 48 hours followed by intracellular flow cytometry for RORγt and IL17A (F) Tamoxifen-treated *Stat3*^fl/fl^ and *Stat3*^fl/fl^CD4^CreER(T2)^ mice were subjected bile duct ligation (BDL) or sham surgery followed by cytometric enumeration of liver infiltrating immune cells, qPCR of whole liver mRNA, and colorimetric assay for serum ALT five days later (**G-I**). Liver sections were stained with H&E for subsequent image analysis of area of necrosis (**J**). Dots represent results from individual wells of at least two independent experiments in **A, B, D, and F** from mice in **C, G-J.** One-way ANOVA was applied with Dunnett’s multiple comparisons test against H_2_O in (**E**) or against “TCDCA+DMSO” in (**F**). Unpaired t-tests were applied for comparisons between two groups except for Mann-Whitney test in (**I/J**) with *p <0.05 and **p<0.01. See also Figure S5.

### STAT3 controls Th17 polarization of hepatic Tregs under cholestatic conditions

In Tregs of patients with latent autoimmune diabetes, phosphorylated (p)STAT3 was reported to bind to the promoter of *FOXP3*, and STAT3 activation was linked to progressive decline in this cell population^47^. We evaluated STAT3 as a candidate regulator of *Foxp3* in Tregs under cholestatic conditions. Indeed, *Stat3* was induced and phospho-STAT3 level was elevated upon TCDCA stimulation (Fig 4A, Fig S5A). In vitro stimulation of splenic Tregs exposed to TCDCA upregulated IL17A production and increased co-expression of RORγt and pSTAT3 (Fig 4B). Upon stimulation with CDCA, splenic Tregs upregulated pSTAT3, which was blocked with S1PR1 but not S1PR4 antagonist (Fig S4E). MDR2^-/-^ mice with chronic cholestasis and liver injury had an increased number of liver infiltrating IL17A+CD25+CD4+ cells which expressed higher levels of pSTAT3 compared with hepatic CD4+ cells from WT mice (Fig 4C, Fig S5B). IL17A (GFP) expression was increased in splenic naïve CD4 cells from IL17A-GFP reporter mice when cultured under Treg differentiating conditions in presence of TCDCA (Fig S4F). In circulating Tregs from healthy blood donors, TCDCA induced signature genes of Th17 polarization, including *LIF* and *IL17F* (Fig S4G). To directly test the role of STAT3 in controlling the balance of Treg/TH17 polarization of CD4 cells, STAT3 activity was blocked with S31-201 which partly prevented TCDCA induced repression of FOXP3 and fully abrogated upregulation of RORγt and IL17A expression in Tregs in vitro (Fig 4E/F). Splenic Tregs from transgenic *Stat3*^fl/fl^CD4^CreERT2^ mice, in which treatment with tamoxifen deleted *Stat3* in CD4 lymphocytes, displayed higher FOXP3 expression after TCDCA exposure in vitro than Tregs from control mice with intact STAT3 signaling (Fig S5D). Deletion of *Stat3* in lymphocytes reduced contraction of the hepatic Treg compartment during acute cholestasis from DDC injury and attenuated hepatocellular injury in this model, as measured by serum ALT (Fig S5E/F). Retention of BA is the primary cause of injury in the BDL model of SC, unlike the DDC model in which precipitation of porphyrin crystals leads to bile duct epithelial and hepatocellular damage and secondary cholestasis. When BDL was performed in *Stat3*^fl/fl^CD4^CreERT2^ mice following tamoxifen exposure, acute cholestasis induced reduction of hepatic Tregs was almost completely prevented (Fig 4F/G). This was accompanied by decreased liver infiltration by CD8 lymphocytes, upregulated whole liver *Areg* and diminished proinflammatory *Tnfa* and *Ifng* mRNA expressions, lower serum ALT levels, and decreased hepatocyte necrosis compared with control mice (Fig 4H-J, Fig S5G).

### Anticholestatic therapy restores hepatic Treg responses in sclerosing cholangitis

We reasoned that if BA suppress hepatic Treg responses in SC, a reduction of hepatic BA concentration ought to restore protective Treg responses. Experiments were designed based on prior studies which showed that treatment with SC-435, a minimally absorbed IBAT inhibitor (IBATi), increased fecal excretion of BA, reduced total liver and serum BA concentrations by 65% and 99%, respectively, and halted progression of SC in MDR2^-/-^ mice ^31^. In MDR2^-/-^/*Foxp3*(GFP) reporter mice, reduction of BA pool size following SC-435 treatment was accompanied by significant increase in frequency and number of hepatic Tregs, decreased IL17A expression by hepatic Tregs, and concomitant decrease in circulating levels of proinflammatory cytokines (Fig 5A/B; Fig S6A/B). There was no evidence of an increase in gut derived Tregs, based on similar expression of CD103 and Neuropilin by hepatic Tregs from both treatment groups (Fig S6C). To explore the role of Tregs in mediating the hepatoprotective effects of IBATi, we treated MDR2^-/-^ mice with SC-435 and simultaneously depleted Tregs with an CD25-antibody (Fig S6D). Compared with untreated MDR2^-/-^ mice serving as controls, SC-435 treatment increased Tregs and decreased CD8 cell only in mice receiving IgG, not those undergoing Treg depletion with αCD25 (Fig 5C). While liver and serum BA concentrations and serum TB levels were significantly reduced in mice treated with SC-435 irrespective of Treg depletion, serum ALT and ALP reduction was abrogated in Treg depleted mice compared to mice treated with SC-435 and IgG (Fig 5D/E). Treg depletion also prevented the histopathological improvement observed in MDR2^-/-^ mice receiving SC-435 and IgG, including reduction in periportal inflammation, ductal proliferation, hepatocyte necrosis, and fibrosis, as assessed by scoring of liver histopathology on a scale of 0 to 4+ on H&E- and SR-stained liver sections (Fig 5F). The semiquantitative evaluation was corroborated by image analysis of SR and CK19+ stained areas for percent fibrosis and intrahepatic biliary mass, respectively (Fig 6F/G). There was no evidence of liver injury in non-cholestatic MDR2^+/-^ mice exposed to SC-435 treatment and Treg depletion (Fig S6E-J). In mice fed chow admixed with 0.1% of DDC and 0.008% of SC-435 for 14 days, treatment with IBATi prevented cholestasis induced reduction of hepatic Tregs and decreased cholestasis, CD8 cells, and injury to the bile duct epithelium and hepatocytes, as shown by improved serum TB, ALP, ALT levels and bile acids (Fig S7A/B). This was accompanied by diminished periportal inflammation, fibrosis, and intrahepatic biliary mass, as demonstrated by image analysis of SR staining and CK19 IHC of liver sections (Fig S7C-E).

**Figure 5.**
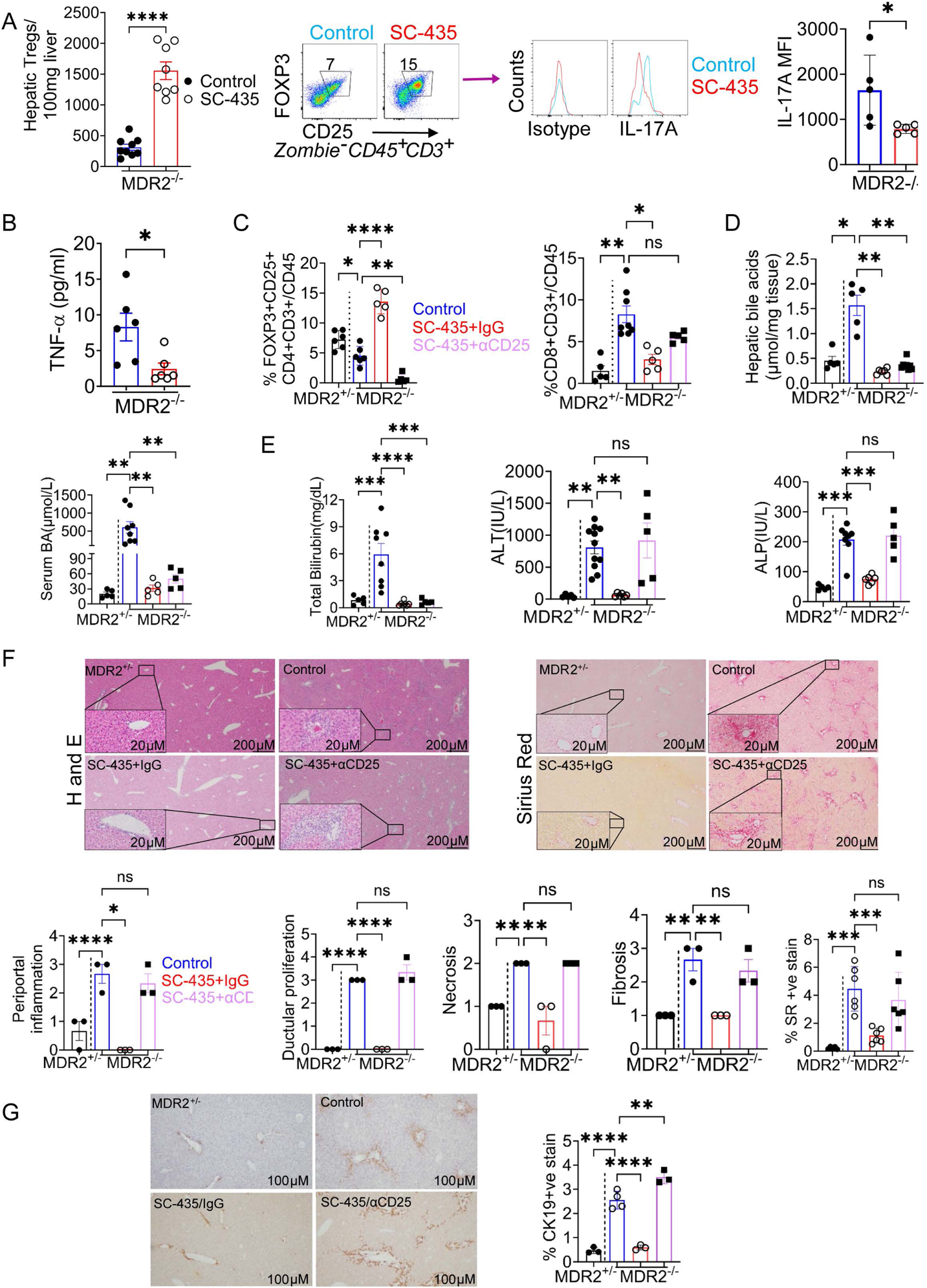
Disruption of intestinal bile acid reclamation promotes infiltration of the liver with Tregs. FOXP3 (EGFP)/MDR2^-/-^ double transgenic mice were fed chow containing 0.008% of the IBAT inhibitor SC-435 for 14 days. Hepatic Tregs were enumerated in freshly isolated liver mononuclear cells and expression of IL17A in Tregs was assessed by flow cytometry following PMA/Ionomycin restimulation (**A**). End of treatment plasma samples were subjected to Luminex assays (**B**). MDR2^-/-^ mice were assigned to three treatment groups: control (blue), SC-435 and IgG (red), or SC-435 and αCD25 to deplete Tregs (purple). Untreated MDR2^+/-^ mice served as non-cholestatic controls (black). Frequencies of intrahepatic Tregs and CD8 lymphocytes at the end of treatment were determined by flow cytometry (**C**). End of treatment hepatic and serum concentrations of total BA, and serum levels of biochemistries, including total bilirubin, ALT, and ALP, were measured with colorimetric assays (**D/E)**. Liver sections were stained with H&E and Sirius Red, and severity of portal inflammation, ductal proliferation, hepatic necrosis, and fibrosis were scored on a 0 to 4+ scale. Percent fibrosis was determined by image analysis (**F**). Liver sections were subjected to CK19 IHC and image analysis for quantification of intrahepatic biliary mass (**G**). Each dot represents an individual mouse. Unpaired t test (in **A/B**) and one-way ANOVA with Dunnett multiple comparisons in MDR2-/- mice against the group “MDR2^-/-^ control” were applied to test for statistical differences between groups with ^ns^non-significant, *p <0.05; **p<0.01, ***p<0.001, and ****p<0.0001. See also Figures S6 and S7.

**Figure 6.**
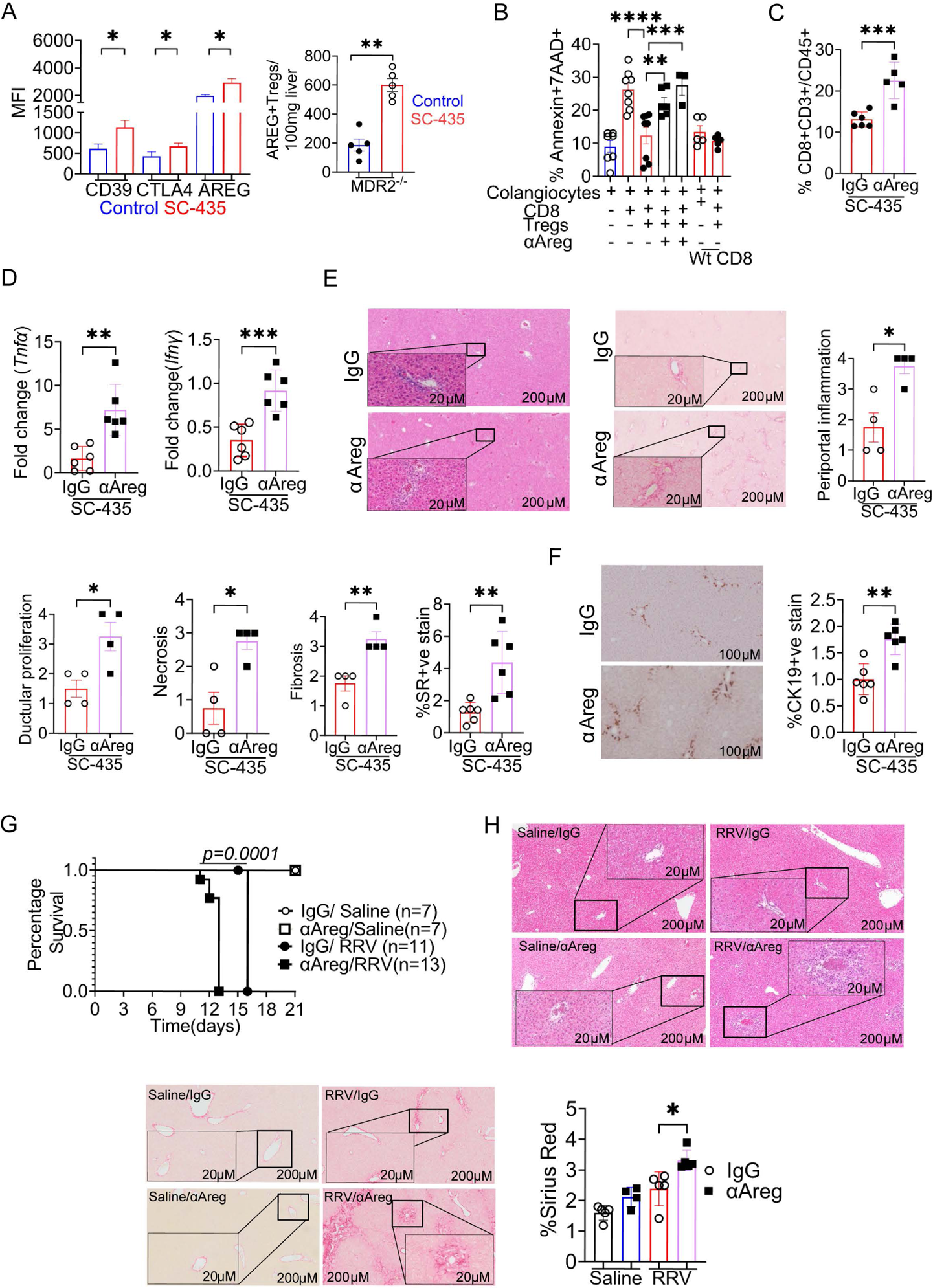
Protection of cholangiocytes from immune mediated injury by Tregs is mediated by amphiregulin. FOXP3 (EGFP)/MDR2^-/-^ double transgenic mice were fed chow containing SC-435 (or without SC-435 in controls) for 14 days prior to cytometric determination of expression of suppressor molecules on Tregs (**A**). Neonatal cholangiocytes were cultured for 24 hours in presence of hepatic CD8 cells from MDR2^-/-^ mice and hepatic Tregs from WT mice in presence of AREG neutralizing antibody (1 or 5μg) prior to cytometric determination of apoptotic cholangiocyte death by 7AAD and annexin staining. Cholangiocytes co-cultured with hepatic CD8 cells from WT mice served as controls (**B**). MDR2^-/-^ mice received AREG neutralizing antibody (or IgG in controls) and SC-435 for 14 days prior to cytometric enumeration of hepatic CD8 lymphocytes **(C**), quantitation of proinflammatory cytokine production by whole liver qPCR (**D**), scoring of liver histopathology on H&E and Sirius Red (SR) stained liver sections (**E**), image analysis of percent liver fibrosis on SR stained sections, and determination of intrahepatic biliary mass by CK19 IHC (**F**). Neonatal mice were injected with saline (n=7 pups (open circles/squares) or rhesus rotavirus (RRV) (n=11-13 pups (closed circles/squares) within 24 hours after birth and subsequently treated with AREG neutralizing antibody (or IgG in controls). Survival was monitored for 21 days (**G**). Liver samples were collected from non-infected mice and from moribund RRV infected mice over time and subjected to H&E and SR staining and image analysis (**H**). Dots represent results from individual mice in all panels except for **C** in which they represent wells of two independent experiments. The unpaired t test was applied to test for statistical significance in difference between groups except for **B** with one-way ANOVA with Dunnett’s multiple comparison test with *p <0.05; **p<0.01, ***p<0.001, and ****p<0.0001. Log rank was used to compare the Kaplan Meyer curves in **G**. See also Figure S8.

### Treg-derived amphiregulin constrains immune mediated bile duct epithelial injury

We examined whether treatment with IBATi restored expression of inhibitory molecules on Tregs previously shown to be downregulated by TCDCA. IBATi-associated reduced cholestasis was linked with upregulated expression of the suppressor molecules CD39, CTLA4, and AREG on hepatic Tregs in MDR2^-/-^ mice and to lesser extent in the DDC model (Fig S8A, Fig 6A, Fig S7F). To investigate the role of Treg-derived AREG in controlling immune mediated bile duct epithelial injury, we co-cultured neonatal mouse cholangiocytes with WT hepatic Tregs and hepatic CD8 lymphocytes from MDR2^-/-^ mice with or without AREG neutralizing antibody (Fig 6B, Fig S8B). Apoptotic death of cholangiocytes cultured with CD8 cells, measured by 7AAD and annexin-5 positive staining, was reduced in the presence of Tregs but was abrogated in dose-dependent fashion when media was conditioned with AREG neutralizing antibodies. To assess the protective role of AREG in vivo, AREG signaling was blocked during IBATi treatment of MDR2^-/-^ mice. Compared with MDR2^-/-^ mice receiving SC-435 and IgG, treatment with SC-435 and neutralizing anti-AREG antibody increased liver infiltration with CD8 lymphocytes, upregulated whole liver pro-inflammatory cytokine expression, worsened the scores of SC histopathology, and increased percent liver fibrosis and intrahepatic biliary mass, as assessed by image analysis of SR staining and CK19 IHC, respectively (Fig S8C/D, Fig 6C-F). These findings were corroborated by upregulation of pro-fibrogenic gene expression in cholestatic mice, in which AREG was neutralized during IBATi treatment (Fig S8E). Mice treated with recombinant AREG before and during DDC challenge displayed an attenuated SC phenotype with lower serum liver biochemistries, increased Treg liver infiltration, and lower number of total and CD44+ CD8 cells, compared with mice receiving PBS and DDC (Fig S8F-I). We then examined the role of AREG in the established model of rhesus rotavirus (RRV) induced neonatal bile duct obstruction ^15, 48^. AREG neutralization shortened survival following post-natal infection with RRV, worsened liver inflammation, and accelerated liver fibrosis (Fig 6G/H).

### Upregulated hepatic Treg- and *AREG* responses at diagnosis confer better surgical outcomes in infants with extrahepatic biliary atresia

In patients with EHBA, pSTAT3 expression was elevated in circulating Tregs compared with age matched healthy controls (Fig 7A). Accumulation of pSTAT3+CD4+ lymphocytes was detected by multiparameter immunofluorescence (IF) in liver biopsies from infants with EHBA (Fig 7B). The multi-center Childhood Liver Disease Research Network (ChiLDreN) has collected biospecimen and prospective clinical data from more than 900 infants enrolled at the time of diagnosis of EHBA. A gene signature derived from bulk RNA sequencing (RNAseq) of diagnostic liver biopsy and persistent elevation in serum BA levels after surgery were reported to predict 2yrSNL ^49, 50^. In order to examine whether liver infiltration with Tregs at the time of diagnosis of EHBA is linked to surgical outcomes, we derived a gene set predictive of hepatic Treg responses in 40 infants with EHBA enrolled into ChiLDreN who had both liver RNAseq data and formalin-fixed paraffin embedded (FFPE) liver sections from diagnostic liver biopsies. Clinical data from these study participants is included in Table S4. Liver sections were subjected to IF and subsequent automated image analysis using antibodies for CD4, CD8, FOXP3, RORγt, TBET, and panCK (Fig 7C). Based on the average count of FOXP3+CD4+ Tregs per area, subjects were divided into groups with low or high Treg counts. A set of differentially expressed genes from these 40 subjects was derived from corresponding liver RNAseq, which was available through the GEO database (GSE122340) (Table S5) ^49^. Differentially expressed genes (DEGs) included several genes which were also upregulated in Tregs compared with other CD4 subsets in the DDC models of SC, including *CCL20, IL1RL1, IL6, KLF4,* and *AREG* (Fig 7D). The DEGs were subsequently applied to RNAseq data from diagnostic liver biopsies of a cohort of 130 infants with EHBA and 10 with idiopathic neonatal cholestasis enrolled into the ChiLDreN PROBE study. 126 pathways of the Gene Ontology (GO) resource were upregulated in cluster A vs B, including three linked to immune function (Fig S9A, Table S6). Pathways of IL-10 signaling, and IL-8 production were upregulated in patients with increased Tregs, and pathways of positive regulation of lymphocyte regulation were correlated with *AREG* gene expression (Fig S9B/C). As expected, of the 40 patients with IF and RNAseq data, subjects in cluster A displayed higher counts of liver infiltrating FOXP3+CD4+, but also of RORγt+CD4+ cells (Fig 7E, Fig S9D). No significant differences for counts of CD8+ or TBET+ cells were revealed between the clusters (Suppl Fig 9E). Considering the cohort of 118 EHBA patients with liver RNAseq and prospectively collected clinical data, including 2-year-outcomes and central review of diagnostic liver histopathology, patients in cluster A had better 2yrSNL and displayed less severe periportal inflammation at diagnosis (Fig 7F/G). No significant differences were found among age, serum liver biochemistries at diagnosis, fibrosis at diagnosis, or bilirubin at 3 or 6 months after HPE (Fig 7H, Fig S9G/H). Considering the 34 patients with liver sections for IF and information on 2ySNL, we found lower number of CD8 lymphocytes and higher ratio of Treg/CD8 cells in patients with better surgical outcomes (Fig 7I). The association between abundance of hepatic Tregs and surgical outcomes in EHBA supports the protective role of Tregs in progression of fibroinflammatory liver disease.

**Figure 7.**
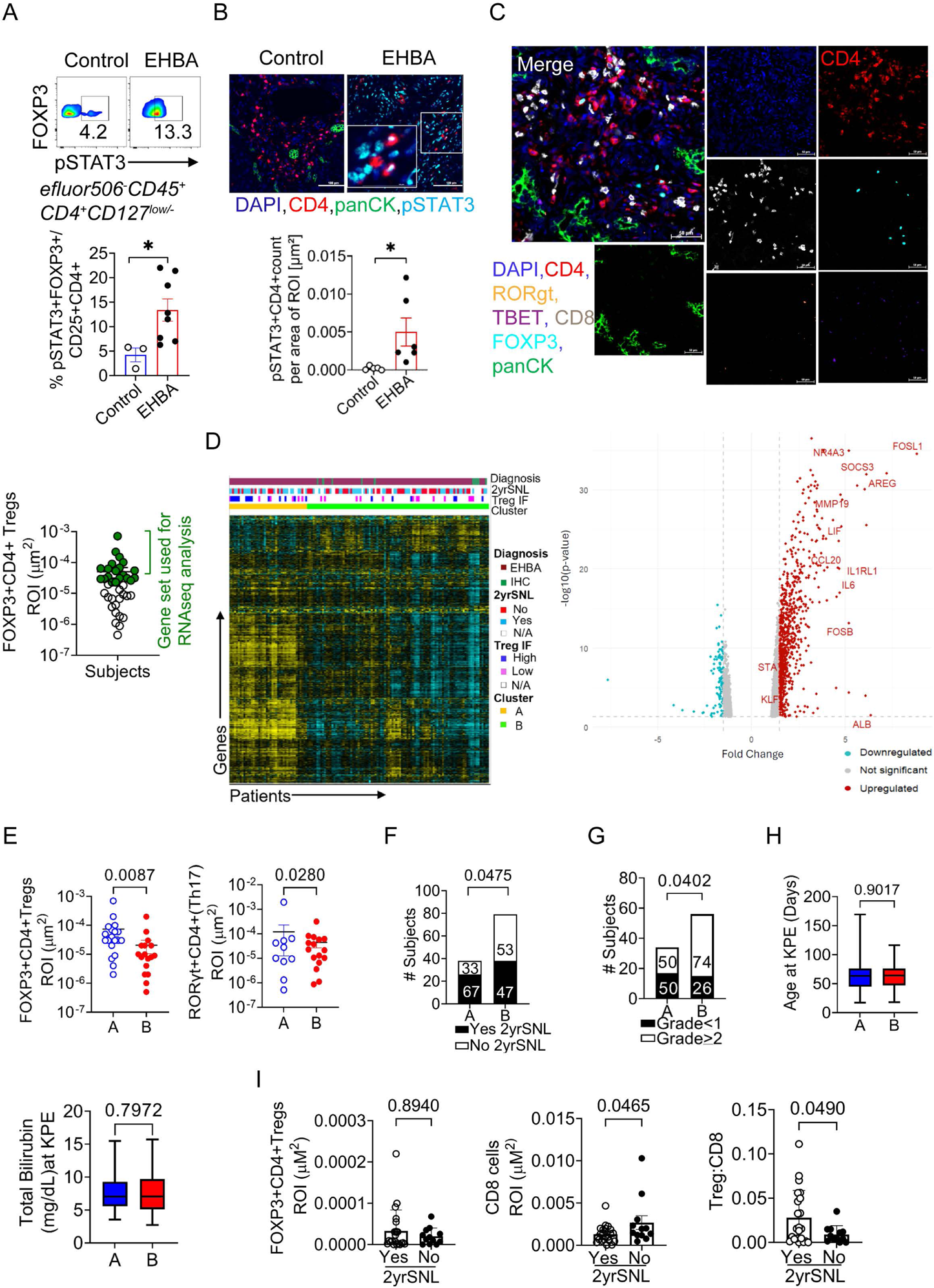
Abundance of hepatic Tregs at diagnosis of extrahepatic biliary atresia (EHBA) is associated with better surgical outcome of hepatoportoenterostomy. Cryopreserved PBMC from patients with EHBA or healthy infants were subjected to intracellular flow cytometry (**A**). Representative photomicrographs of multi-parameter IF on liver sections from patients with EHBA at diagnosis and non-cholestatic age matched controls. CD4+pSTAT3+ cells were enumerated as average # cells/portal tract (**B**). Representative images of multi-parameter IF on a liver biopsy from an infant with EHBA. Number of FOXP3+CD4+Tregs per area ROI was determined by automated image analysis in a cohort of 40 patients with EHBA. Subjects with number of Tregs in top 50% percentile (highlighted in green) were designated as “high” Tregs (**C**). A set of differentially expressed genes was derived from liver RNAseq data of subjects with “high” vs “low” Treg counts by IF and applied to the entire cohort of infants with liver RNAseq data available from the time of diagnosis with either EHBA (n=130) or idiopathic neonatal cholestasis (INH; n=10). Unsupervised analysis identified 2 clusters of patients based on similar expressions of Treg associated genes. Volcano plot displays the upregulated genes in cluster A vs B with log FC>1.5 and p-value <0.05 in red (**D**). For 40 patients with IF and automated image analysis on liver sections, the numbers for liver infiltrating FOXP3+CD4+ and RORgt+CD4+ lymphocytes are plotted according to their assignment to liver RNAseq based clusters A or B (**E**). Survival outcomes at 2 years (2yrSNL, **F**), grades of portal inflammation as scored by central review of liver histopathology (**G**), age and serum liver biochemistries were plotted based on cluster assignment by liver RNAseq (**H**). Frequency of liver infiltrating immune cells determined by IF was compared between patients stratified by 2ySNL (**I**). Each dot represents an individual patient. Unpaired t test and Fisher exact tests were applied to test for significant differences between groups. See also Figure S9 and Tables S4-6.

## Discussion

The experimental data on mechanisms by which the liver microenvironment shapes homeostasis and function of hepatic Tregs under cholestatic conditions is relevant for several liver diseases, such as EHBA, PSC, PBC, or chronic liver allograft rejection. While primary BA and their metabolites have been studied extensively for their effects on Tregs and Th17 lymphocytes in the intestine, this is the first study to our knowledge investigating the effects of conjugated BA on Tregs in the liver ^51–54^. We demonstrate that TCDCA suppresses Treg-maintenance and functions involving DNMT3-mediated hypermethylation of *Foxp3* and RORγt and STAT3-dependent upregulation of Th17-associated genes. Interventions to block these pathways or reduce hepatic retention of conjugated BA boost the number and protective function of hepatic Tregs, diminish hepatic CD8 responses, and attenuate hepatobiliary injury in complementary mouse models of SC. Our findings expand on prior reports in preclinical models of cholestatic liver disease, indicating a critical role for Tregs in constraining bile duct epithelial injury and subsequent biliary fibrosis ^6, 18–20, 55, 56^. We link Tregs to cholestatic liver disease phenotype in humans by showing that preserved hepatic Treg response in a subset of infants with EHBA is associated with better 2yrSNL.

To delineate the impact of cholestasis on Tregs, we surveyed hepatic T lymphocyte transcriptional profiles in the DDC model of SC with time course resolution. Using scRNAseq and ATACseq, we present a regulatory atlas of 16 cell clusters encompassing epithelial, endothelial, and immune cells relevant for hepatobiliary injury and repair in SC. We describe three CD4 memory T-cell populations, including Treg, which are distinguished by mRNA expression and chromatin accessibility of *Foxp3, Ctla4*, and *Areg*.

Severe cholestasis between D7 and D14 of DDC challenge is accompanied by induction of gene regulatory networks in Tregs involving the TFs FOS, FOSB and RORC as regulators of Th17-associated genes *Il17a* and *Il23r*. FOS has been implicated in Th17 polarization in models of atopic dermatitis and multiple sclerosis ^57^. We confirm the predictions from single-cell (sc)-genomics studies by demonstrating increased IL17A, RORγt, and STAT3 expression in Tregs cultured in the presence of TCDCA and in hepatic Tregs purified from cholestatic MDR2^-/-^ mice. The propensity of CD4 lymphocytes to adopt a Th17 phenotype was recently described in patients with PSC in single cell resolution ^58^. *In vitro*, hydrophobic BA repress suppressor function of natural Tregs without reducing their viability or inducing apoptosis and reduce induction of Tregs from naïve CD4 cells. The latter process is likely the primary cause for diminished number of hepatic Tregs observed in the DDC, BDL and MDR2^-/-^ models of SC. Cholestasis induced decrease in hepatic Treg number is accompanied by rise in hepatic CD8 lymphocytes. Vice versa, restoration of suppressor phenotype in natural Tregs and surge in hepatic Treg-number from induced Tregs inhibit CD8 lymphocyte responses following treatment of cholestatic mice with IBATi. Probing the molecular mechanisms, we show that TCDCA directly affects polarization of Tregs by hypermethylating *Foxp3* promoter elements, and upregulating *Stat3*. Inhibition of DNMT by 5-Aza prevents *Foxp3* hypermethylation and attenuates the SC phenotype in MDR2^-/-^ mice. While hypermethylation of *Foxp3* in Tregs had not previously been attributed to direct effects of BA, experiments in cholestatic neonatal mice from RRV-induced EHBA showed a surge of hepatic Tregs following treatment with 5-Aza ^59^. Genetic deletion of *Stat3* in CD4 lymphocytes partly protect Tregs from TCDCA-induced repression of FOXP3 in vitro, but more prominently preserves the number of hepatic Tregs in SC, especially after BDL, suggesting a negative effect of STAT3 activation on Treg induction under cholestatic conditions. The findings are consistent with recent observations in which treatment with the STAT3 antagonist shifted the balance of Tregs/Th17 cells in experimental encephalomyelitis ^60, 61^ and disrupted STAT3 signaling CD4 lymphocytes promotes Treg responses and reduces lethality in murine acute graft versus host disease ^62^. While there is paucity of literature on the effects of BA on hepatic CD4 lymphocyte homeostasis, primary BA were shown to control CD4 lymphocyte function in the small intestine regulated by nuclear receptor CAR and the export pump MDR1, preventing BA toxicity ^41,42^.

Our sc genomics studies predict S1P receptors to mediate conjugated BA effect on Tregs and not canonical BA receptors like FXR or TGR5. Indeed, Tregs from FXR-/- or TGR5-/- mice are not protected from TCDCA induced FOXP3 repression. In contrast, pharmacological inhibition of S1PR1 prevents repression of FOXP3 and induction of RORgT and pSTAT3 in purified Tregs stimulated and cultured in presence of CDCA or TCDCA in dose dependent fashion. Our conclusion that S1PR mediates the effects of TCDCA on hepatic Treg function is supported by reports on the role of S1PR controlling macrophage activation and lymphocyte polarization. For instance, secondary BA like DCA activate macrophages following binding to S1PR2 and aggravate experimental colitis^63^. Its conjugate GDCA upregulates ZBP1 in macrophages promoting necroptosis and accelerated fibrosis in a BDL model with relevance for EHBA ^64^. Importantly, activation of S1PR by its endogenous ligand sphingosine 1 is linked to STAT3 activation in Tregs, their infiltration into tumor tissue ^65^, and Th17/Treg balance in rheumatic heart disease and aplastic anemia^66, 67^.

Our investigations on the effects of IBATi on hepatic Treg homeostasis in the MDR2^-/-^ model of SC corroborate the effects of BA on Tregs in vitro and suggests a new mechanism of action by which this class of anticholestatic drugs might impact cholestatic liver disease progression. Treatment of MDR2^-/-^ mice with IBATi blocking intestinal BA reclamation lowers serum and liver BA concentration and improves the SC phenotype ^31^, but also promotes liver infiltration with AREG-expressing Tregs. Protection from progression of SC is abrogated when Tregs are depleted, or AREG is neutralized. A surge in number of hepatic Tregs is accompanied by diminished bile duct epithelial and liver fibrosis following treatment with IBATi is validated in the DDC model of SC. Our experiments suggest that restoration of hepatic AREG/Treg responses may contribute to the mechanisms of action by which IBATi treatment improves cholestatic liver disease. The potential immunomodulatory effects of IBATi have not been examined in clinical trials in patients with cholestatic liver disease from Alagille syndrome, PFIC, PBC, or PSC ^68–70^. A clinical trial is currently investigating the impact of IBATi on 2ySNL in infants with EHBA (ClinicalTrials.gov identifier: NCT04336722). It will be important to ascertain in clinical studies of IBATi in cholestatic liver disease how hepatic Treg responses are influenced by treatment and whether restoration of hepatic Treg responses is linked to treatment effects on liver clinical endpoints. Our findings also emphasize the importance of the liver microenvironment in the design of clinical trials on low dose IL-2 immunotherapy to promote Tregs in AIH and PSC ^71^. Under severe cholestatic conditions such treatment may need to be combined with anticholestatic therapies to enhance protective Treg responses at the site of tissue damage.

The hallmark of EHBA is severe cholestasis at diagnosis. While the functions of CD8, Th17, and Treg lymphocyte populations as effector and regulatory cells are well established in mouse models of the disease, there is only limited evidence that these processes operate in infants with the disease^7, 12, 18, 19^. Here we begin to validate our findings from animal models in experiments utilizing human samples. For instance, TCDCA induces a gene signature of Th17 polarization in purified Tregs from healthy blood donors, pSTAT3 is abundantly expressed by liver CD4+ and circulating Tregs from infants with EHBA, and gene expression signatures of preserved Treg and *AREG* responses are associated with decreased periportal inflammation and improved 2yrSNL in infants with EHBA.

Increased infiltration of the liver with FOXP3+Tregs but also with RORgt+CD4+ cells is observed in patients with better surgical outcomes. It is currently unknown whether the RORgt+CD4+ cells are derived from Tregs and whether these cells contribute to disease progression. Our group showed before that IL17A+ CD4+ lymphocytes orchestrate recruitment of inflammatory macrophages to the biliary epithelium in experimental EHBA and prevalence of IL17A+ cells is correlated with infiltration of the portal tracts with CD68+ macrophages in infants with EHBA ^7^. Collectively, our results suggest that AREG-expressing Tregs are critical for protecting cholangiocytes from immune mediated injury, and their function is determined by conjugated BA retained in the liver under cholestatic conditions. The impact of anticholestatic therapies on restoring these protective immune responses and the relationship with liver disease endpoints requires carefully designed clinical trials in EHBA and PSC.

## Acknowledgements

The authors would like to thank Dr. Matthew Kofron, the director of the Confocal Imaging Core at the Cincinnati Children’s Hospital for the help with confocal imaging and image analysis. The authors would also extend their gratitude to patients and families for participating in the observational studies, hepatobiliary surgeons Drs. Bondoc and Mullapudi and clinical research coordinators E. Chapman and J. Hawkins for facilitating liver tissue sample collection. We appreciate the assistance by Drs. Kostrub and Vig from Mirum Pharmaceutical in providing the IBAT inhibitor, SC-435, on the basis of MTA.

## Author contributions

RK, performed the animal-based experiments and RK, PB, LP, AT, MM, CSL, SP, MS and AYvH performed biochemical assays and histological staining. IO and TS performed the multiparameter IF. HJ performed the pyrosequencing assay. RK performed the bulk RNAseq analysis. AB reviewed liver histopathology. DAH, GT, CC, WZ contributed experimental design and provided human tissue samples. ZFY, JAW, ERM and AGM designed and ZFY performed the sc-genomics analyses. SS, SH and NS aided in liver cell type annotation of sc-genomics data. AGM designed the experiments. All authors contributed to drafts of the manuscript.

## STAR Methods

### RESOURCE AVAILABILITY

#### Lead contact

Further information and requests for resources and reagents should be directed to the lead contact, Alexander Miethke.

#### Materials Availability

This study did not generate new unique reagents.

#### Data and code availability

Raw and processed snRNA-seq and snATAC-seq have been deposited in GEO under accession numbers GSE230085 and GSE230091, respectively. Human hepatic mRNA levels were analyzed in gene expression data originally obtained from bulk hepatic RNAseq studies carried out on patients with EHBA enrolled into PROBE (n=140) ^72^. Detailed information on handling of liver biopsy samples, RNAseq protocols, internal controls, normalization procedures, analysis of gene expression, and corresponding outcome data were deposited in GEO (GSE122340) ^73^. This paper does not report original code. Any additional information required to analyze the data in this paper is available from the lead contact upon request.

### EXPERIMENTAL MODEL AND STUDY PARTICIPANT DETAILS

#### Human Studies

##### Patient Information

Unstained sections from liver biopsies or explants, peripheral blood mononuclear cells (PBMC), and clinical data were obtained from patients enrolled into the Prospective Database of Infants with Cholestasis (PROBE; NCT00061828), a protocol of the National Institute of Diabetes and Digestive and Kidney Diseases–supported Childhood Liver Disease Research Network (ChiLDReN), or a prospective cohort study at Cincinnati Children’s Hospital Medical Center (CCHMC; Hepatic Immune Activation: #20123320). The studies were approved by the ChiLDreN ancillary study committee and the Institutional Review Board at Cincinnati Children’s Hospital Medical Center (CCHMC) and conformed to the ethical guidelines of the 1975 Declaration of Helsinki. Informed consent was obtained from parents/guardians. Liver tissue as collected at the time of diagnostic liver biopsy or HPE. Diagnosis of EHBA was confirmed by review of liver histopathology and intrahepatic cholangiograms confirming large duct obstruction. Demographics, disease status, and serum biochemistries at the time of collection of PBMC for flow cytometric studies are listed in Table S7.

#### Murine Studies

C57BL6/J wildtype (WT), BALB/cJ WT, C.Cg-Foxp3^tm2Tch^/J (Foxp3^EGFP^), and B6(129X1)-Tg(Cd4-cre/ERT2)11Gnri/J (CD4CreER^T2^) mice were purchased from Jackson Laboratory. BALB/cAnNCrl (BALB/cCR) mice were purchased from Charles River. MDR^2*-*^*^/-^* mice in the BALB/cJ background were a gift from Dr. Frank Lammert (Homburg University; Homburg, Germany) ^29^. *Nr1h4* (FXR) deficient C57BL/6J mice (*Nr1h4*^-/-^) were a gift from Dr. Frank Gonzalez (National Cancer Institute, NIH) ^74^. *Gpbar1 (*TGR5) deficient 129S3/SvImJ×C57BL/6 mice (*Gpbar1^-/-^)* were gifts from Dr. Galya Vassileva (Merck Research Laboratories)^75^. IL-17^GFP^ mice were received from Biocytogen LLC (Worcester, MA) through MTA and backcrossed into the BALB/cJ strain at CCHMC animal facility for 10 generations ^7^. Double transgenic MDR2^-/-^/*Foxp3*^EGFP^ mice were generated in-house by breeding MDR2^-/-^ mice with Foxp3^EGFP^. David Hildeman gifted us with B6.129S1-Stat3^tm1Xyfu^/J (Stat3^fl/fl^) from Jackson Laboratory as well as transgenic mice with inducible *Stat3* in CD4 lymphocytes, generated in his lab by breeding Stat3^fl/fl^ with CD4CreER^T2^ to produce Stat3^fl/fl^CD4CreER^T2^ ^76^. All age and sex information are included in the relevant methods section. Mice were housed at Cincinnati Children’s Hospital Medical Center (Cincinnati, OH). Mice were bred in-house and group-housed in a pathogen-free animal facility at 25°C, 12-hour light/dark cycle, with ad libitum autoclaved chow, unless otherwise stated. All experiments were performed with the approval of the Institutional Animal Care and Use Committee of Cincinnati Hospital Medical Center and in accordance with US Department of Health and Human Services, Office of Laboratory Animal Welfare and Association for Assessment and Accreditation of Laboratory Animal Care (AAALAC) International guidelines.

### METHOD DETAILS

#### Immunofluorescent staining

Multiplex-IF was applied to FFPE tissues using Opal™ 7 color solid Tumor Immunology Kit (PerkinElmer). On the same tissue section, pan-cytokeratin (panCK 1:750, Abcam), TBET (1:750, Abcam) CD8 (undiluted, Ventana), CD4 (1:1250, Cell Marque), FOXP3 (1:50, Abcam), RORgT (1:1000, Abcam) and DAPI were detected. Following deparaffinization, slides were rinsed for 5 min with deionized water and placed in a plastic container filled with antigen retrieval (AR) buffer that was preboiled for 1 min via microwave at 100% power, until bubbles formed. Slides were then microwaved for 15 min at 20% power and cooled for 30 min at RT, rinsed in deionized water, then rinsed in 1X Tris-buffered saline with 1% Tween-20 (TBST). Tissue sections were blocked for 10 min and incubated with primary antibody for either 1h at RT or overnight at 4°C depending on the antibody. Slides were incubated for 10 min with opal polymer HRP Ms + Rb secondary antibody followed by 10 min Opal fluorophore incubation then mounted with ProLong Gold antifade reagent with DAPI (Invitrogen).

##### Image acquisition and automated analysis

Slides were imaged using Inverted, motorized Nikon Ti-2 widefield microscope. Tile scan slide images taken with a 20X objective and subjected to blind spectral unmixing. Marker expressions were automatically quantified using NIS-Element image analysis software (Nikon). Cells were quantified and normalized to tissue area.

#### In vitro experiments with FACS purified Tregs

To determine the effects of BA on human Tregs, PBMCs were isolated from the buffy coat of healthy, adult blood donors using Ficoll (Thermo Fisher Scientific) density gradient method. Tregs staining 7AAD^-^ CD3^+^CD4^+^CD127^-/low^CD25^+^ cells were purified by FACS (SH800, Sony) and collected in 1ml of 100% FBS. FACS sorted cells were cultured with α-CD3, α-CD28 (Beads 1:1 ratio), hIL-2 (100U), TCDCA (100 μM), or vehicle (H_2_O) in prime -XV T cell expansion media (Irvine Scientific) for 48 hours prior to downstream assays including flow cytometry or RNAseq.

For RNAseq, cells were washed and centrifuged at 12000 rpm for 2 minutes at 4°C prior to snap freezing of the pellets for RNA isolation. Cells were lysed with lysis/binding buffer and total RNA was extracted according to the manufacturer’s protocol and eluted in 60µl Elution Buffer (mirVana miRNA Isolation Kit). RNA integrity was determined by Bioanalyzer (Agilent, Santa Clara, CA). Libraries for RNA-seq were prepared by using NEBNext Ultra II Directional RNA Library Prep kit (New England BioLabs) at the University of Cincinnati Sequencing Center (med.uc.edu/departments/eh/cores/genomics). To study differential gene expression, individually indexed and compatible libraries were proportionally pooled (∼25 million reads per sample in general) for clustering in cBot system (Illumina, San Diego, CA). Libraries at the final concentration of 15 pM were clustered onto a single read (SR) flow cell v3 using Illumina TruSeq SR Cluster kit v3 and sequenced to 51 bp using TruSeq SBS kit v3 on Illumina HiSeq system. For analysis, sequence reads were aligned to the genome using standard Illumina sequence analysis pipeline, which was analyzed by the Laboratory for Statistical Genomics and Systems Biology at the University of Cincinnati.

#### Flow cytometric studies

PBMCs were isolated from EDTA blood samples of infants with EHBA and healthy infants using Ficoll density (Thermo Fisher Scientific) gradient centrifugation and cryopreserved until flow cytometric assays were performed in batches.

##### Surface staining

Cells were stained with fixable viability dye (Thermo Fisher Scientific) and incubated with anti-CD16/32 FC block (BioLegend) or mouse serum (Thermo Fisher Scientific) for 15 min at 4°C, washed with FACS buffer, spun at 2000rpm at 4°C for 2 min, resuspended and incubated with surface antibodies for 20 min at 4°C prior to intracellular staining or acquisition on cytometer.

##### Intracellular/nuclear flow cytometry

For intranuclear staining of FOXP3 or pSTAT3, cells were fixed with fixation/permeabilization buffer (ThermoFisher Scientific) or with BD Cytofix/Cytoperm (BD Biosciences), respectively, according to the manufacturers’ protocols. For FOXP3 staining, cells were fixed by adding 50ul of fixation buffer (Thermo Fisher Scientific), incubated at room temperature for 20 min, washed using 1X permeabilization buffer (Thermo Fisher Scientific), followed by centrifugation at 2000 rpm for 2 min. Cells were then incubated with FOXP3 antibody diluted in permeabilization buffer for 30 min at 4°C. Cells were stained for pSTAT3 (BD Biosciences) using True-Phos perm buffer in cell suspensions protocol (Biolegend). Cells were washed and acquired on a SORP LSRII or Fortessa (BD Pharmingen). Flow cytometric analysis was performed using FlowJo (Tree star). Down sample, TSNE and flowsom plugins were used to generate TSNE plots of cytometry data.

#### Hepatic RNAseq studies in infants with EHBA

Of the 140 patients with RNAseq and clinical data, we received sections of formalin fixed paraffin embedded (FFPE) biopsy samples from 40 patients at the time of diagnosis of EHBA. These sections were subjected to multi-parameter IF to enumerate hepatic Tregs. Corresponding hepatic gene expression data was interrogated to determine differential gene expression between patients with more than the median number of Tregs of the entire cohort. The differential gene signature was subsequently applied to the entire cohort of 140 patients with EHBA and single end (se)RNAseq data ^49^ and clusters of patients with similar gene expression were identified by unsupervised analysis. To derive a Treg-associated gene set all reasonably expressed genes (TPM >3 in >20% of the total cohort; n=12903 genes) were included in the analysis, which used a moderated *t-*test to identify differentially expressed genes between high/low Treg patients. We identified n=1265 genes with a cutoff of *p*<0.05; further filtration identified 425 genes with fold change >1.5. Hierarchical clustering of all available patients in the EHBA/ICH cohort using the significant genes (Complete linkage rule and Pearson’s Centered similarity measure) identified two overall clusters (assigned ‘A’ and ‘B’). To assess the association of Treg-based clusters with clinical characteristics, we used Fisher’s exact test for categorical variables (2-year survival with native liver and baseline inflammation grading) and t*-*test for continuous variables (liver biochemistries and age).

#### Animal experiments

##### DDC model of Sclerosing Cholangitis

8-week-old, male C57Bl/6J mice were fed with 5058 Picolab Mouse Irradiated chow (Cincinnati Lab Supply, OH) admixed with 0.1% of 3,5-diethoxycarbonyl-1,4-dihydrocollidine (DDC) for up to 14 days^32^[1 and then changed to 5058 control chow for up to 4 weeks to model recovery from hepatobiliary injury. Groups of mice were harvested before (D0), during DDC treatment (D7/14), or after challenge (D14+7/14/28) to obtain liver tissue and plasma samples for downstream assays, including scRNA and snATACseq. To examine the effects of Tregs on sclerosing cholangitis phenotype, mice were injected intra-peritoneally (i.p) with anti-CD25 antibody (250μg/dose; clone PC-61.5.3, BioXcell) or isotype control IgG1 (250μg/dose; HPRN, BioXcell^6^ matched IgG every 48 hours starting at 4 days prior to DDC treatments for 7 days.

To confirm the role of amphiregulin in hepatic protection and repair during cholestasis, we supplemented in vivo amphiregulin levels by administrating 1.5-5µg mouse recombinant amphiregulin (R&D systems) or PBS vehicle (100μL). 8-week-old female BALB/cJ wild-type untreated mice and DDC-treated mice were administered recombinant amphiregulin or vehicle every other day starting 2 days prior to DDC treatment and concluding 7 days post DDC exposure.

To assess the impact of BA on regulatory T cells in DDC model of SC, 8-week-old female BALB/cJ mice were fed with irradiated 5058 diet admixed with 0.1% DDC and 0.008% ASBT inhibitor SC-435 (Mirum Pharmaceutical, CA) for 14 days providing approximately 11 mg/kg/day of the minimally absorbed compound SC-435 ^77^.

To address the role of Stat3 in controlling Treg T cell number and function 6-8-week-old, male Stat3^fl/fl^CD4^CreER(T2)+/-^ were treated daily with 20mg of tamoxifen via IP injection for 4 days prior to administering 14 days of 0.1% DDC admixed chow or 5058 control chow.

##### MDR2-/- Model of Sclerosing Cholangitis

To determine how cholestasis impacts Treg homeostasis, 30-day-old, female Mdr2^-/-^ mice were administered 20 µg/g body weight of 5-Aza daily for seven days via i.p. injection. In studies examining the effects of BA reduction on hepatic Treg homeostasis, 30-day-old, female Mdr2^-/-^ mice were fed irradiated 5058 chow admixed with 0.008% SC-435 for 14 days ^31^. To deplete Tregs or neutralize AREG during SC-435 treatment, 250 µg of aCD25 antibody (clone PC-61.5.3, BioXcell) or 5 µg of anti-AREG antibody (R&D systems)^78^ were administered via i.p. every 48 hours for 18 days starting 4 days prior to beginning SC-435 treatment. Control mice received isotype matched antibodies; Treg depleted mice received HPRN monoclonal rat IgG1 (BioXcell) or polyclonal goat IgG (Bioxcell) for AREG neutralization during SC-435 treatment.

##### Bile duct ligation model of sclerosing cholangitis

6-8-week-old, male Stat3^fl/fl^CD4^CreER(T2)+/-^ were treated with 20mg of tamoxifen daily via i.p for 4 days prior to bile duct ligation (BDL). Stat3^fl/fl^ mice without cre served as controls. While under anesthesia, mice underwent upper-midline laparotomy, the common bile duct was ligated with coated VICRYL (polyglactin 910) 3-0 suture, and skin and fascia were closed with absorbable suture. In sham treated control mice, laparotomy was performed without ligation of the bile duct. Livers and serum samples were collected for downstream assays 5 days after BDL.

##### Rhesus rotavirus-induced experimental EHBA

Neonatal BALB/cCR mice were injected s.c. with 1.5 x10^6^ ffu (focus forming units) of rhesus rotavirus (RRV)^19^ within 24 hours of birth to induce experimental biliary atresia, as previously described ^18^. To examine the role of AREG in EHBA, 2µg/mouse/day of anti-AREG antibody (or isotypic IgG as control) in PBS was administered on days 3,5,7 post-RRV infection and mice were monitored for survival for 21 days. Liver tissue samples were collected from euthanized moribund mice and subjected to sirius red staining and image analysis.

##### Measurement of serum biochemistries

Blood samples collected by cardiocentesis at the time of harvest were spun at 2300 rpm for 20 minutes at 4°C and subjected to colorimetric assays to measure alanine aminotransferase (ALTCatachem), alkaline phosphatase (ALPCatachem\), or total bilirubin (TBPointe Scientific) with Chemistry Calibrator Set (Pointe Scientific). Samples were measured for absorbance according to manufacturer protocols using Biotek Synergy H1 microplate reader ^31^.

##### Measurement of liver bile acid concentrations

Liver tissue (50-100mg) was homogenized at room temperature in 1 mL of 75% ethanol using an Omni Bead Ruptor 4, followed by incubation for 2 hrs at 50°C, and centrifugation for 10 min at 6000 xg at 4°C. The supernatant was collected for bile acid analysis using the mouse Total Bile Acids Kit (Crystal Chem) and absorbance was measured using Biotek Synergy H1 microplate reader at 540 nm.

##### Sclerosing cholangitis histology assessment

H&E and Sirius red staining of liver sections was performed according to standard procedures and severity of periportal inflammation, ductal proliferation, necrosis, fibrosis was scored on a 1-4+ scale by pathologist A.B. who was blinded to mouse genotype and treatment assignment, as previously validated^6, 79^.

##### CK19 immunohistochemistry

Formalin-fixed liver sections were subjected to dewaxing, deparaffinization, and antigen retrieval with 1X AR pH9 buffer (AR9). Further quenching of endogenous peroxidase with 3% H2O2 was followed by blocking with 5% Normal Goat Serum (Jackson) in PBS/1%BSA, incubation overnight at 4°C with Troma III (1:100) primary antibody (Developmental Studies Hybridoma Bank, University of Iowa), then washed and incubated with goat anti-rat secondary antibody. Slides were washed and incubated with Vectastain Elite ABC-HRP Reagent (Vector Laboratories) followed by DAB Peroxidase (Vector Laboratories) staining for visualization ^6^.

##### Image analysis

Image analysis of CK19 and Sirius Red stained liver sections was performed by digital scanning of liver sections at 10x using a Nikon H550S (Nikon Instruments, Inc, Tokyo, Japan). Scanned slides were analyzed by fraction of DAB positive area or collagen staining in 15 randomly selected 1000um×1000um sections per liver sample using NIS-Elements software (Nikon Instruments, Inc, Tokyo, Japan) ^6^.

#### Protein and gene expression quantification

##### Western blot

Protein from Tregs, non-Treg CD4 cells, and liver was extracted using IP lysis buffer (ThermoFisher Scientific) with cOmplete ULTRA (Roche) and phosSTOP (Sigma Aldrich). Concentration of extracted proteins was measured using the Pierce BCA kit (ThermoFisher Scientific). Protein samples were boiled in Laemmli buffer prior to separation by SDS-PAGE (Invitrogen), transferred onto polyvinyl difluoride (PVDF) membrane (Bio Rad), and incubated with primary antibody overnight at 4°C. Primary antibodies were diluted in BSA and included rabbit anti-mouse DNMT3a, pSTAT3 and alpha tubulin (Cell signaling). Blots were washed and incubated with Donkey anti-rabbit IR Dye 800CW secondary antibody (Li-Cor Biosciences) for 2 hrs at room temperature. Li-Cor Biosciences Odyssey Classic Imager was used to visualize immunoblots.

##### Luminex assay

TNFα and IL6 concentrations were measured in 25 µl of serum by ProcartaPlex multiplex immunoassay kit (Thermo Fisher Scientific) at the CCHMC Flow Cytometry Core according to manufacturer’s instructions ^80^.

##### Real Time PCR

Total RNA was isolated from cultured Tregs using the RNeasy micro-Kit (Qiagen). cDNA was prepared using RevertAid RT Reverse Transcription Kit (ThermoFisher Scientific). qPCR was performed using PowerUp™ SYBR™ Green Master Mix or TaqMan™ Universal PCR Master Mix (ThermoFisher Scientific) on 7900HT PCR system (Applied Biosystems) with gene specific primers or TaqMan probes, and analysis was performed applying the comparative delta CT method. Primers for *18s, FoxP3, Dnmt1, Dnmt3a, Stat3, Col4A3, Col1a2* are listed below in Table 2.

#### Functional assays

##### Isolation of lymphocytes populations

10-day-old newborn and female, 8-week-old Foxp3^EGFP^, BALB/cJ WT, C57BL/6J WT, Nr1h4^-/-^, and Gpbr1^-/-^ mice were used for experiments examining the effects of BA on Treg molecular phenotype and function in vitro. Spleens were harvested, minced, and incubated with Collagenase D for 15 min at 37°C with 5% CO_2_ then deactivation with FBS, passed through 70µm nylon mesh, and washed after centrifugation at 300 g at 4°C for 7 min. Following red blood cell lysis, cells were washed, and the pellet was resuspending in MACS buffer. CD4^+^CD25^+^ Tregs were separated using Miltenyi beads (Miltenyi Biotech) according to manufacturer’s instructions. For some experiments, Tregs were FACS purified on Sony cell sorter (Sony) as efluor506-CD45^+^CD3^+^CD4^+^CD25^+^GFP^+^ cells following incubation with the respective antibodies in serum coated tubes filled with 1 mL of 100% FBS. CD8 T lymphocytes were separated from splenocytes or liver mononuclear cells using the Dynabeads™ FlowComp™ Mouse CD8 Kit according to the manufacturer’s protocol^6^.

Naive non-Tregs from the liver and spleen were purified from 8-week-old females BALB/cJ WT mice by sorting CD62L+CD44-CD4+ T cells and cultured in presence of human IL-2 (100U/mL), dynabead Mouse T-Activator CD3/CD28 (Bead to cell ratio 1:1), and TGF-β1 (10 ng/mL) for 72 hrs with hydrophobic bile acids TCDCA and CDCA.CD45RA positive naive CD4+ CD8-cells were isolated according to manufacturer protocol (Miltenyi). Naive CD4 cells were cultured according to the same protocol as the naive non-Tregs.For intracellular detection of IL17A, cells were cultured for 4 hrs in the presence of Phorbol 12-myristate 13-acetate (PMA) (500 ng/mL), ionomycin (500ng/mL) and treated with 0.5 µL of Golgiplug Protein Transport Inhibitor containing Brefeldin A to enhance the detectability of cytokine-production in splenic and hepatic Tregs.

##### Annexin V Staining

To measure percentage of apoptosis in Tregs cultured in presence of CDCA/TCDCA cells were stained with 5 µL Annexin V staining buffer and 10 µL of 7-AAD and incubated for 15 min at RT in the dark. Before flow cytometry 400 µL of Annexin V Binding Buffer was added to each sample.

##### Cell proliferation Assay

Cell proliferation in Tregs cultured with either TCA/TCDCA was measured by using CellTrace Violet Cell Proliferation Kit per manufacturer protocol (ThermoFisher Scientific).

##### In vitro assays with Tregs

MACS sorted splenic Tregs were cultured with αCD3/CD28 (2μg/mL), IL-2 (100U) and BA (CDCA, CA, TCDCA, TCA at concentration as indicated in Figures or legends) or with vehicle (DMSO/dH_2_O for unconjugated/conjugated BA) for 48 hrs. Small molecules to block S1PR1 (FTY720), S1PR4 (CYM50358), STAT3 (S31-201), or methylation (5-aza-2’-deoxycytidine (5-Aza)) were added to the culture media in concentrations as outlined in the respective figure legends. For proliferation assays, Tregs were washed thoroughly after incubation with BA to remove BA and co-cultured with splenic CD8+ cells in a 1:1 ratio with 50,000 cells per well in the presence of αCD3/CD28 (2μg/mL), IL-2 (100U), and 10% FBS for 3 days. CD8 proliferation was measured by Cell Trace Violet dye dilution using flow cytometry.

##### Cholangiocyte apoptosis assay

Cholangiocytes were isolated from 2-day-old BALB/cJ mice and cultured as described previously ^81^. Hepatic Tregs and CD8 cells were isolated from WT and from MDR2^-/-^ mice in BALB/cJ background, respectively, and co-cultured in 1:1 ratio with cholangiocytes in presence of αCD3/CD28 (2μg/mL), IL-2 (100U), and 1 or 5 µg of amphiregulin neutralizing antibody for 24 hours followed by cytometry for 7AAD and Annexin 5 on CK19+ cholangiocytes as indicators of apoptosis ^82^. In brief, cells were washed with MACS buffer after co-culture and resuspended in 100 µL of Annexin V Binding Buffer. 5 µL of FITC Annexin V and 5 µL of 7-AAD viability staining solution were added to each sample and incubated in the dark for 15 min at RT prior to addition of 400 µL of Annexin V Binding Buffer. Cells were washed and acquired on a Fortessa, and analysis was performed using FlowJo.

##### Bisulfite pyrosequencing assay

MACS sorted splenic CD4^+^CD25^+^ natural Tregs from Foxp3^EGFP^ reporter mice were cultured in presence of TCDCA, (5, 100 μM), or TCA, (50, 1000 μM), or with vehicle (dH_2_O) for 48 hrs in presence of αCD3/CD28 (2μg/mL) and interleukin (IL)-2 (100U). Cells were washed and genomic DNA was isolated using the EZ DNA methylation-Gold Kit (Zymo Research). 200 ng of genomic DNA was subjected to sodium bisulfite treatment according to the manufacturer’s protocol (Zymo Research). Standard bisulfite PCR amplification reaction was performed to amplify *Foxp3* gene fragments at an annealing temperature of 50°C. Pyrosequencing was carried out using Pyro Gold reagents with a PyroMark vacuum prep workstation and a PyroMark Q96 MD instrument (Qiagen). The generated pyrograms were automatically analyzed using PyroMark analysis software (Qiagen). The pyrosequencing assay was validated using SssI-treated human genomic DNA as a 100% methylation control and human genomic DNA amplified by GenomePlex^®^ Complete WGA kit (Sigma) as 0% methylation control. **Sequences** for primers to amplify the promoter of *Foxp3* are included in Table S7.

#### Murine single cell genomics studies

##### Isolation of liver mononuclear cells

Liver mononuclear cell (LMNC) suspensions enriched in immune cells were prepared from mice treated with DDC at various timepoint using a gradient centrifugation method ^83^. Briefly, livers were perfused in situ using RPMI media containing 1 mg/mL Collagenase D (Sigma-Aldrich), collected in digestion media (RPMI with 1 mg/mL collagenase D (Sigma-Aldrich) and 10 µg/mL DNase I (Roche)), minced into minute pieces and digested *ex vivo* in 10 mL digestion media, shaking at a 250 rpm at 37°C for 15–20 min. To deactivate Collagenase D, 5mL of heat inactivated FBS was added. Cells were isolated by pushing digested tissue through 100μm strainer. LMNCs were then separated using either 33% Percoll (GE Healthcare) (for control mice) or by using double gradient method (for DDC treated mice), as described previously ^83, 84^. For scRNAseq, 1×10^6^ cells/mL were submitted to the sequencing core, of which 16×10^3^ cells were used for library preparation and sequencing.

##### Stellate, endothelial, and cholangiocyte isolation

For stellate, endothelial and cholangiocyte isolation, mice were perfused with HBSS buffer without Ca^2+^ using a 60 mL syringe. This was followed by perfusion using HBSS with Ca^2+^ containing protease (14 mg/mouse) (Sigma-Aldrich) and HBSS with Ca^2+^ and Collagenase D (20 mg/mouse) via cannulation of the portal vein. All buffers were kept at 37°C and at a pH of 7.4. Following perfusion, livers were collected, minced, and digested using protease (5mg/mL), collagenase (1 mg/mL), and DNase I (10 μg/mL) at 37 °C for 15-30 min at 250 RPM/minute depending on level of liver fibrosis. Following digestion, the cell suspensions were passed through 100µM nylon mesh and divided for specific preparations for stellate cells, cholangiocytes and liver sinusoidal endothelial cells (LSECs). For hepatic stellate cell isolation, cells were resuspended in 8 mL of GBSS buffer and mixed with 4 mL Nycodenz (ProGen) solution with 1.5 mL overlaying with GBSS solution (Sigma-Aldrich). Stellate cells were further separated by density gradient centrifugation as described previously ^85, 86^. Cholangiocytes were purified by gradient centrifugation followed by anti-EpCAM antibody mediated positive selection, as described previously ^81, 82^. LSECs were isolated using Anti-CD146 rat monoclonal (PE (Phycoerythrin) antibody using the EasySep Release Mouse PE Positive Selection Kit (STEMCELL Technologies), per manufacturer’s instructions.

##### Isolation of nuclei for scATAC seq

Nuclei were isolated according to the protocol for isolating liver LMNCs for single cell or single nuclei sequencing ^83^. To separate nuclei from cells, 100 μL of cold Lysis Buffer (10 mM Tris-HCl, 10 mM NaCl, 3mM MgCl_2_, 1% BSA, 0.1% Tween-20, 0.01% Digitonin (ThermoFischer Scientific), 0.1% Nonidet P40 (Sigma-Aldrich)) was added to 1×10^6^ cells and mixed up to ten times with a pipette for up to 120 seconds to lyse the cells. For instance, the best nuclei were obtained from LMNC after 90 sec whereas epithelial cells were only lysed for 60 sec. After cell lysis, nuclei were washed with 1 mL of cold wash buffer (10 mM Tris-HCl, 10 mM NaCl, 3mM MgCl_2_, 1% BSA, 0.1% Tween-20) prior to resuspension in 1 mL of cold 1X Nuclei Buffer (10x Genomics) at a concentration of 1×10^6^ nuclei/mL. The integrity of nuclei was assessed by trypan blue using a hemocytometer. For scATAC, 1×10^6^/mL nuclei were submitted to the CCHMC sequencing core, of which 16 x 10^3^ were used for library preparation and sequencing.

##### Library preparation

Illumina^TM^ system and scRNAseq chromium Next GEM Single Cell 3ʹ Reagent Kits v3.1 (10x Genomics) and scATACseq Chromium Next GEM Single Cell ATAC Reagent Kits v1.1 (10x Genomics) were used by the CCHMC Gene Expression Core for library preparations. Libraries generated from each individual sample were sequenced on a Novaseq 6000 in the CCHMC DNA Sequencing Core.

#### Single-cell RNA-seq data processing

##### Bioinformatic processing and quality control

The raw data was processed through Cellranger 3.1.0 (10X Genomics) for alignment, initial filtering, and feature-barcode matrix generation using mouse reference genome mm10[ftp://ftp.ensembl.org/pub/release93/gtf/mus_musculus/Mus_musculus.GRCm38.93.gtf.gz]. Background noise including random barcode swapping and ambient RNA was removed from the count matrix using CellBender^87^.

Based on a benchmarking paper^88^ comparing computational doublet detection methods, five best-performing algorithms were chosen, including hybrid,^89^ scDblFinder,^90^, DoubletFinder ^91^, Solo^92^ and Doublet Detection (Gayoso & Shor, 2022 https://zenodo.org/record/6349517#.ZFYj6nZBxPY), to predict doublet scores for cells in each sample with default settings. Cells within each sample were ranked by each method alone from high to low doublet score, and ranks were averaged. Cells of high doublet ranks were removed proportionally to the number of doublets expected in each sample, given number of cells recovered and the estimated multiplet rate: 0.8% per 1,000 recovered cells^93^. Downstream analysis was done in R version 4.0.2, following standard workflows from Seurat v4.0.4 ^94^. Cells were further filtered according to following the criteria: total UMI counts between 500 and 40,000, number of genes detected between 200 and 7,000, <15% transcripts mapped to mitochondrial genes, <40% transcripts mapped to ribosomes, and <0.75% of UMI mapped to cell cycle genes. Any genes detected in fewer than 20 cells were removed. After QC filtering, 101,044 quality cells were kept for downstream analysis.

##### Sample integration and cell type annotations

Global-scaling normalization *LogNormalize* (Seuart *NormalizeData* function) was next applied and the top 2,000 most variable genes were selected after variance-stabilization transformation (*vst* method in Seurat *FindVariableFeatures* function). We then mean-centered and normalized by high variable gene variance while regressing out variation due to UMI counts (Seurat *ScaleData* function) and principal component analysis (PCA) was performed for dimension reduction with the top 50 components (Seurat *RunPCA* function). Shared nearest neighbor (SNN) graphs for 10 nearest neighbors with top 50 components (Seurat *FindNeighbors* function) were constructed using Louvain clustering to iteratively group cells at the resolutions from 0.1 to 1.0 (Seurat *FindClusters* function) and visualized in UMAP with non-linear dimensional reduction. Reference mapping was conducted by Seurat anchor-based integration from published liver scRNA-seq datasets^95, 96^ and further refined by marker gene annotation. Using SCTransform normalization^97^, we merged the 19 scRNA-seq samples using 5,000 variable features (Seurat *SelectIntegrationFeatures* function), performed PCA reduction on the combined data and integrated with Harmony 0.1 using the top 50 principal components^98^. After Harmony integration, an SNN graph was built with Louvain clustering and visualized in UMAP. The label transfer prediction scores were visualized per clusters, and majority voting was used to derive consensus cell type annotations for the 16 major populations across resolutions: 0.3,0.5,0.6 and 0.8.

##### Cell subtype annotations of ILC, T cells

22,768 ILC and T cells were extracted, performing label transfer per sample, using log-normalized counts, published liver datasets processed with only ILC and T cell subsets^58,96^, and sample merging based on 2,000 variable features (Seurat *SelectIntegrationFeatures* function). As above, PCA reduction was performed across datasets and integrated with Harmony (top 20 principal components. After harmony integration, SNN graphs were built with Louvain clustering and visualized in UMAP. ILC and T cell subtypes were annotated based on the cell type-specific gene expression of literature-curated markers. Consistent with previous studies ^99–101^, gamma-delta T-cells did not cluster distinctly from ILC and alpha-beta T cells based on scRNA-seq, so this population was defined based on the following rules: *Cd3d*+ and either *Trdc*>*Trac* or *Trdc*>*Trbc*. Given gene dropout^102^, these rules conservatively identified gamma-delta T cells. By scATAC-seq (see below), gamma-delta T cells clustered distinctly, confirming their presence in the data and highlighting the importance of multiome-seq or paired 5’scRNA-seq with sc-TCR-seq for future work.

##### Differential gene expression analysis

To improve reproducibility across biological replicates, differential gene expression analysis was performed using pseudobulk approaches^103^. “Pseudobulk” gene expression profiles were constructed by tracks by aggregating expression across single cells by cell type, time point and biological replicates (n=3 for D0, n=2 for D14, n=3 for D14+14). DEseq2^104^, with a design formula *gene expression ∼ cell type*, was used for differential analysis and data normalization (variance-stabilization transformation (VST)). Cell type-specific gene signatures, used later for GRN analysis, were identified as highly expressed genes with Log2FC>1 and FDR =10% in one cell type relative to at least another cell type but not less expressed with Log2FC<-1 and FDR=10% in any other cell type^105^. Time-dependent genes were identified using the one-factor design formula *gene expression ∼ cell_type_time_point*, with log2|FC|>1, FDR=10%, comparing timepoints within each cell type to each other. Gene expression was visualized with ComplexHeatmap ^106^.

#### Single-cell ATAC-seq processing

##### Bioinformatic processing and quality control

Sequencing data were processed using Cell Ranger ATAC 1.2.0 (10X Genomics) for initial barcode filtering, cut sites identification, and peak calling, using the mouse reference genome mm10[https://cf.10xgenomics.com/supp/cell-atac/refdata-cellranger-arc-mm10-2019-A-1.2.0.tar.gz]. A Seurat object was built with a peak counts matrix and fragment files through Signac^107^ and high-quality cells were kept: peak region fragments between 1,000 and 40,000, >40% fragments in peaks, <3% ENCDOE blacklist^108^ region ratio and a maximum mononucleosomal/nucleosome-free ratio of 2 (Signac *NucleosomeSignal* function). Only peaks on autosomal and X chromosomes were kept. To identify doublets, ArchR^109^ (ArchR *addDoubletScore* function) was used with default parameters. As for scRNA-seq, doublets were removed according to the estimated multiplet rate of 0.8% per 1,000 cells recovered ^42, 93^. After QC filtering, 41,387 quality cells were kept.

##### Peak calling, sample integration, and clusters

To combine all samples, the new reference peak sets were generated across samples using MACS2 peak-calling (version 2.1.4)^110, 111^. The fragment files were first modified per sample to split transposase cut sites with separate entries for both 3’ and 5’ of the fragments and cut sites were filtered out within ENCODE blacklist regions ^108, 112^. Frequency-inverse document frequency (TF-IDF) normalization (Signac *RunTFIDF* function) was then conducted, top variable peak features with min. cutoff=‘q0’ were identified, latent semantic indexing (LSI) dimension reduction (Signac *RunSVD* function) was performed, and the top 50 singular values were computed. Further, graph-based clustering was performed and non-linear dimension reduction by UMAP (Seurat *FindNeighbors* and *FindClusters* functions) was done at the clustering resolution of 0.5. Only LSI components that have less than 0.8 Pearson correlation with total cut sites in peaks were used. Small clusters with <200 cells were merged to the most similar larger clusters based on Spearman correlation analysis of DEseq2 VST normalized pseudobulk accessibility per cluster, then peaks were called within each sample by merged clusters using MACS2, with extension=50, shift=-25, effective genome size=2.6e9, FDR=0.05. The common sets of reference peaks across samples were merged via intersection (GenomicRanges package *reduce* function ^113^), and only peaks with a cut site in at least 100 cells per sample were retained. This process resulted in 330,504 reference peaks. Fragments within reference peak regions were re-quantified for each sample to get the new peak fragment count matrix, then merged and integrated scATAC-seq samples using Seurat anchor-based method on the LSI embeddings (Signac *IntegrateEmbeddings* function). Finally, cells were clustered across all samples on the integrated LSI embedding through the Louvain algorithm as above, considering resolutions 0.1 to 0.9.

##### scRNA-seq label transfer for cell major types

To define the cell types in scATAC-seq gene expression was imputed from scATAC-seq based on the number of cut sites mapped to coordinated gene promoter regions with 2kb upstream and 500bp downstream extension of TSS (Signac *GeneActivity* function), then normalized by total cut sites within each cell, scaled by median counts across cells, and log-transformed. After PCA dimension reduction, label transfer was perfomed from scRNA-seq to scATAC-seq estimated gene activity through Seurat canonical correlation analysis and anchors were identified using the top 50 principal components (Signac *FindTransferAnchors* and *TransferData* functions). Cell types were defined by majority voting for each cluster based on label transfer predicted cell types at the resolution 0.9, as determined from clustree 0.5.0 analysis ^114^.

##### ILC, T cell subtype characterization

To define ILC and T cell subtypes for scATAC-seq, reference peaks were built specifically for these cell types. Separately for 13,330 ILC and T cells, dimension reduction was perfomed within each sample, cell clusters were identified based on Louvain clustering at 0.5 resolution using LSI components < 0.8 correlation to cut sites per cell. Small clusters (<200 cells) were merged to the closest large cluster based on correlation analysis. Peaks were called through MACS2 per sample using the following parameters: extension=50, shift=-25, effective genome size=2.6e9, and 5% FDR cutoff. Common sets of peaks across samples were used for the new count matrix for each sample and then merged for the integrated ILC-T cell object. Further integration with ILC and T cell-only scRNA-seq object were conducted using imputed gene activities within each sample. The predicted labels were added to the integrated ILC and T cell scATAC-seq object, and cell types were determined through majority voting at cluster resolution 1.0.

##### Differential accessible peaks analysis

Mirroring our scRNA-seq analysis, ATAC-seq signal was aggregated by cell type and time point into pseudobulk signal tracks and differential chromatin accessibility analysis was perfomed and normalized using Deseq2 (design formula: *accessibility ∼ cell type* and variance-stabilization transformation (VST)). Cell type-specific peaks were identified as highly accessible regions (Log2FC>1 and FDR=10% in one cell type relative at another cell type) and not less accessible (Log2FC<-1 and FDR=10% relative to another cell type. Results were visualized using ComplexHeatmap.

#### CD4 T cells gene regulatory network inference

##### Target gene selection from RNA-seq

To understand dynamic gene regulation in Tregs following DDC-induced liver cholestasis, gene regulatory networks (GRNs) were built for encompassing the memory CD4+ T cell populations. The subset of memory CD4+ T cells scRNA-seq data was log normalized and *vst* transformed. Gene models were constructed for the top 2,000 most variable genes in the dataset.

##### Potential regulators selection

Potential regulators were selected based on the following criteria: 1) TFs within the curated mouse TFs list collected in ^41^ as targets (top 2000 genes with variable expression); 2) Essential regulators in Th17 response based on literatures^115, 116^. A total of 177 TFs (Table S3. Potential regulators) were identified as potential regulators to build the network.

##### Prior network from scATAC-seq

A prior network of TF-target gene interactions was constructed based TF binding site (TFBS) predictions at accessible chromatin regions cis to gene loci^41, 42^. In brief, we generated and merged cell-type specific reference peaks for each memory CD4+ T cell populations, as above, resulting in a total of 96,051 peaks. For TFBS prediction, mouse and human TF binding motifs position weight matrices (PWMs) from the CIS-BP motif 2.0 database^117, 118^ were downloaded in MEME format and split into 100 files for parallel motif scanning. Peaks were scanned for motif occurrences using FIMO^119^ (thresh=0.00001, max-stored-scores=500,000 and a first-order Markov background model.) TFs were associated with genes if the motif occurrence (P_raw_<10^-5^) fell within +/- 10kb of the gene body^41^. The prior network included a total of 59,795 TF-gene interactions between 120 TFs and 1842 genes (Table S3. Prior network).

##### Inferelator framework

The *Inferelator* model ^105^ was applied for GRN inference to single-cell settings, which models gene expression as a sparse multivariate linear combination of transcription factor activity using a modified LASSO -StARs framework as follows:

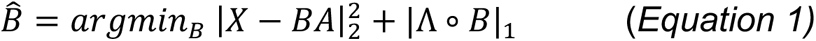

where B ∈ ℝ^|genes|×|TFs|^ is the matrix of inferred TF–gene interaction coefficients, Λ ∈ ℝ^|genes|×|TFs|^ is a matrix of nonnegative penalties, and ° represents an entry-wise matrix product. X ∈ ℝ^|genes|×|samples|^ is the expression matrix for genes in the prior, A ∈ ℝ^|TFs|×|samples|^ is the unknown TFA. The TFA matrix A can further be estimated through either TF gene expression (TF mRNA) or based on the prior network-based TF-gene interactions as follows:

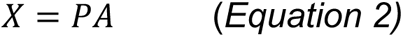

where P ∈ ℝ^|genes|×|TFs|^ is the prior matrix of ATAC-derived TF–gene interactions. Because both TF mRNA and prior-based TFA have advantages, methods were constructed using both approaches.

The LASSO penalty term Λ_ik_ was used to incorporate prior information from scATAC-seq into the network inference using prior network-based TFA estimation, Λ_ik_ bias 0.5 was used to reinforce the TF-gene interactions supported by the prior network. To rank TF-gene interactions by confidence, the stability-based method StARS^120^ was applied, using 500 subsamples of size 880 cells (11.4% of the N=7,746 cells, according to StARS subsample size (N) formula: 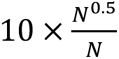 and instability cutoff=0.05. As recently benchmarked on single-cell data^42^, the resulting GRNs from prior-based TFA and TF mRNA modeling were combined by taking the maximum edge confidences up to an average of 10 TFs per gene model, and filtered to retain TF-gene interactions with partial correlation > 0.01. The final GRN contained 19,818 interactions between 134 TFs and 1740 target genes (Table S3. Final network).

##### Treg core TFs, signature genes, and GRNs

To identify candidate cell type-specific core TFs, we started with a low-stringency differential analysis of Treg and CD4 TRMs for TF gene expression at single-cell resolution using the Wilcox test (Seurat *FindMarkers*) and TF activity using a two-sided, unpaired t-test. TFs overexpressed in one cell type relative to another were kept with FDR=10%, and Log2(FC) > 0.25. We took the union of differential TFs from mRNA and TF activity-based tests and ended up with 107 TFs as potential core TFs. To determine core TFs as activators or repressors within each cell type, we performed gene set enrichment analysis (GSEA) for the TF activated and repressed targets from the final network with FDR cutoff 5%, using cell type-specificity up and down-regulated signature gene sets as described previously^41^. Signature genes were defined through DEseq2 analysis of pseudobulk signal tracks, comparing Treg with CD4 TRMs, using the design formula of *gene expression ∼ cell type*, with cutoffs FDR = 10%, Log2FC>1(Table S1. Cell-type specific Treg signature genes; Table S2. Time specific Treg signature genes). Core GRNs for Treg were extracted from the combined network using the GSEA enrichment of Treg-specific gene targets and Treg core TFs, resulting in a final Treg core network with 19 TFs, 1,318 target genes, and 4,812 edges (Table S3. Treg core network).

##### Network visualization

For GRN node visualization, genes were colored according to gene expression and TFs according TF activity estimates. For gene expression, a time point and cell type-specific pseudobulk gene expression matrix was generated, as above, and z-scored across pseudobulk conditions. The pseudobulk TFA matrix was estimated using pseudobulk gene expression and final GRN, according to Equation 2, and z-scored. Treg core GRNs were visualized using jp_gene_viz (https://github.com/simonsfoundation/jp_gene_viz) ^41^. In Figure 2J, visualization was focused on Treg TFs and target genes involved in Th17 polarization and bile acid signaling.

##### Statistical analysis

Graphs were generated using GraphPad Prism version 8 (GraphPad, La Jolla, CA). Values were expressed as mean±SEM unless otherwise specified. *p* values were determined by unpaired, two-tailed *t-*test or by one-way analysis of variance (ANOVA) with Dunnett’s post hoc test for grouped analysis, with multiplicity adjusted *p* values (GraphPad); *p*<0.05 was considered significant. Details on statistical tests and number (n) of data points are provided in the relevant figure legends.

## Abbreviations

2yrSNL: 2-year survival with native liver
5-Aza: 5-aza-2’-deoxycytidine
ATACseq: Assay for Transposase-Accessible Chromatin sequencing
AREG: Amphiregulin
BD: Bile duct ligation BSEP-Bile salt export pump
CA: Cholic acid
CDCA: Chenodeoxycholic acid
ChiLDreN: Childhood Liver Disease Research Network
CTLA4: Cytotoxic T-lymphocyte-associated protein 4
CXCR3/6: C-X-C Motif Chemokine Receptor 3/6
DDC: 3,5-diethoxycarboncyl-1,4-dihydrocollidine
DEG: Differentially expressed genes
DNMT3a: DNA (cytosine-5)-methyltransferase 3A
EGFR: Epidermal growth factor receptor
EHBA: Extrahepatic biliary atresia
FFPE: Formalin fixed paraffin embedded
FOXP3: Forkhead box P3
FXR/NR1HR: Farnesoid X receptor
GPBAR1/TGR5: G protein-coupled bile acid receptor 1
GRNs: Gene regulatory networks
GSEA: Gene set enrichment analysis
HPE: Hepatoportoenterostomy
IBAT: Ileal bile acid transporter
LMNC: Liver mononuclear cell
LSEC: Liver sinusoidal endothelial cells
LSI: Latent semantic indexing
NTCP: Sodium taurocholate co-transporting polypeptide
PSC: Primary sclerosing cholangitis
PWMs: Position weight matrices
RORγt: RAR-related orphan receptor gamma
RRV: Rhesus rotavirus
S1PR1: Sphingosine-1-Phosphate Receptor 1
SC: sclerosing cholangitis
sc-genomics: single cell genomics
scRNA-seq: Single-cell RNA sequencing
TBET: T box transcription factor TBX21
TCDCA: Taurochenodeoxycholic acid
TF: Transcription Factor
TFBS: Transcription Factor binding site
VST: Variance-stabilization transformation

## Supplemental Material

**Table S4:**
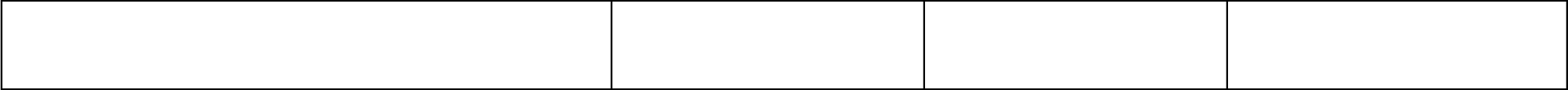
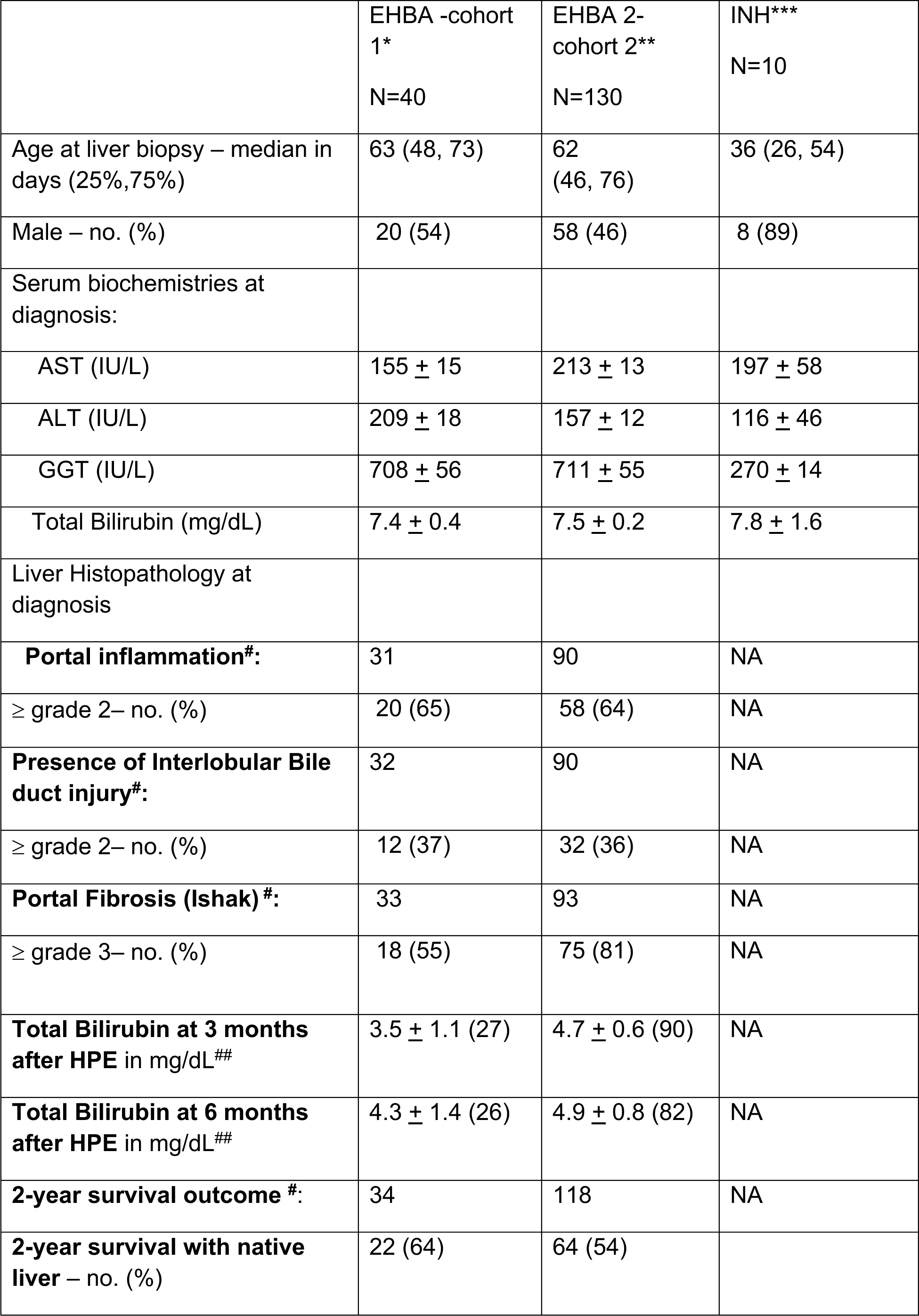
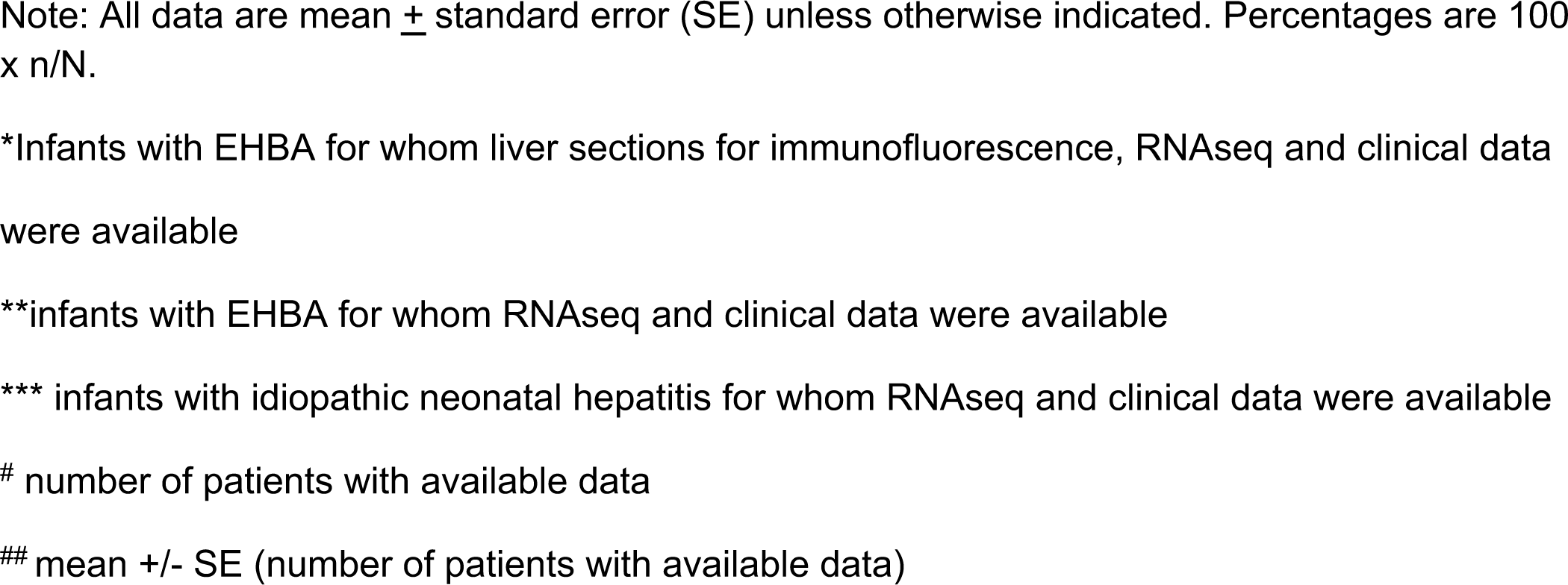
Baseline characteristics and outcomes for infants with cholestasis enrolled into the liver RNAseq study, Related to Figure 7.

|  | EHBA -cohort<br>1* | EHBA 2-cohort 2** | INH***<br>N=10 |
| --- | --- | --- | --- |
|  | N=40 | N=130 |  |
| Age at liver biopsy – median in days (25%,75%) | 63 (48, 73) | 62 (46, 76) | 36 (26, 54) |
| Male – no. (%) | 20 (54) | 58 (46) | 8 (89) |
| Serum biochemistries at diagnosis: |  |  |  |
| AST (IU/L) | 155 ± 15 | 213 ± 13 | 197 ± 58 |
| ALT (IU/L) | 209 ± 18 | 157 ± 12 | 116 ± 46 |
| GGT (IU/L) | 708 ± 56 | 711 ± 55 | 270 ± 14 |
| Total Bilirubin (mg/dL) | 7.4 ± 0.4 | 7.5 ± 0.2 | 7.8 ± 1.6 |
| Liver Histopathology at diagnosis |  |  |  |
| **Portal inflammation#:** | 31 | 90 | NA |
| ≥ grade 2– no. (%) | 20 (65) | 58 (64) | NA |
| **Presence of Interlobular Bile duct injury#:** | 32 | 90 | NA |
| ≥ grade 2– no. (%) | 12 (37) | 32 (36) | NA |
| **Portal Fibrosis (Ishak) #:** | 33 | 93 | NA |
| ≥ grade 3– no. (%) | 18 (55) | 75 (81) | NA |
| **Total Bilirubin at 3 months after HPE in mg/dL<sup>##</sup>** | 3.5 ± 1.1 (27) | 4.7 ± 0.6 (90) | NA |
| **Total Bilirubin at 6 months after HPE in mg/dL<sup>##</sup>** | 4.3 ± 1.4 (26) | 4.9 ± 0.8 (82) | NA |
| **2-year survival outcome #:** | 34 | 118 | NA |
| **2-year survival with native liver – no. (%)** | 22 (64) | 64 (54) |  |

Note: All data are mean ± standard error (SE) unless otherwise indicated. Percentages are 100 x n/N.
\*Infants with EHBA for whom liver sections for immunofluorescence, RNAseq and clinical data were available
\*\*infants with EHBA for whom RNAseq and clinical data were available
\*\*\* infants with idiopathic neonatal hepatitis for whom RNAseq and clinical data were available

**Table S6:** Pathways of differentially regulates genes between cluster A and B in 130 patients with neonatal cholestasis and EHBA, Related to Figure 7.

| S. No | Pathway | q-value<br>FDR B&H | Hits (genes) in Query List |
| --- | --- | --- | --- |
| 1 | Enzyme regulator activity | 1.64x10<sup>-8</sup> | SERPINA3, SPRY2, SPRED2, CRIM1, RPS6KA3, PLEKHG1, DLC1, WFDC2, CDC42EP2, CCL2, CCL3, SIPA1L2, ARHGAP21, ALOX5AP, ARAP2, IPO7, ARHGAP23, RGS5, ANXA1, ANXA2, ANXA2P2, ANXA3, ANXA5, RGCC, BIRC3, CST7, DIS3, AREG, AZIN1, ARHGAP5, GNA13, SOCS1, C9orf72, PHACTR1, ARHGAP42, STARD13, CAB39, ARPP19, SWAP70, TAGAP, PPP1R12B, ARHGAP26, PRSS22, NCF2, BCL2A1, BCL2L1, TNFRSF10D, TNFRSF10B, TNFRSF10A, ANKLE2, IQGAP1, FGL2, TINAGL1, SOCS2, CFLAR, STK4, BMP2, MALT1, BNIP2, GCLM, GMFB, RABGEF1, GNA12, BCL10, C5AR1, DENND5A, DNMBP, KALRN, CALU, MAP3K14, SOCS3, CAST, CAV1, NCS1, TBC1D8, MMP25, CCNH, P2RX7, TIMP3, PPP1R15B, SERPINE1, CD44, PAK2, PIK3R5, CDKN1A, CDKN2B, NRG1, PDGFB, SOCS6, PPP1R15A, KLF4, SH3BP4, SERPINB8, SERPINB9, PLAA, PIM1, AGFG1, TNFAIP8, RHOF, HSP90AA1, SH3BP5, COL6A3, MTMR12, WARS1, PLEKHG2, PPARD, YWHAG, RIN1, PPP2R2A, PPP2R5E, RASGEF1B, ITPRIP, PRKCE, RAPGEF2, MAPK8, ARHGEF26, MAP2K1, DNAJC3, USP6NL, PRNP, RAPGEF5, CDC42SE1, TBC1D5, CD55, LXN, DDX3X, JUN, CCNYL1, PTPRC, ARFGAP3, CCNL1, PZP, PPP4R1, RALGDS, RARRES1, RASA2, RCAN1, TFPI2, HBEGF, ARHGEF5, RGS2, RGS4, RGS16, ABCE1, ARHGAP12, SOCS4, EIF5, MOB1A, |
|  |  |  | SERPINB1, SPRED1, PSME3, RP2, RPGR, TRIB1 |
| 2 | Cell adhesion molecule binding | $3.01 \times 10^{-7}$ | LRRC59, ADAM8, PICALM, EHD4, S100P, ESYT2, FGFR1, TAGLN2, SNX9, SDCBP, SELL, ANXA1, ANXA2, SLC3A2, MSN, MAPRE1, PLPP3, GNA13, PALLD, SWAP70, SPP1, ADAM9, IQGAP1, CCN4, EHD1, ADAM17, CALD1, CAST, THBS1, CD9, PAK2, CDH6, CDH11, PACSIN2, VAPA, NRG1, PFKP, CEMIP2, TJP2, HSPA1A, VASP, CNN2, CNN3, VCL, EZR, DNAJB1, ICAM1, BAG3, CPE, YWHAZ, PPL, CHMP2B, MICALL1, CCN1, CCN2, BZW1, IL1B, SLK, TES, ITGA3, ITGAV, ITGAX, ITGB1, ITGB3, DDX3X, DDX6, PTPN1, F11R, PVR, KIF5B, JAM3, KRT18, DSG2, LAD1, RDX, LCP1, S1PR3, EPHA2, EIF5, LYN |
| 3 | Cytokine binding | $2.07 \times 10^{-5}$ | NBL1, GBP1, GBP2, IL18R1, NRP2, CCRL2, TGFB3, TGFBR2, TGFBR3, THBS1, CD44, IL1RL1, TNFRSF1A, OSMR, TNFRSF1B, CCR1, ZFP36, IFNGR1, CSF2RA, IL1R1, IL1RAP, IL1RN, IL4R, IL6R, IL6ST, ITGAV, ITGB1, ITGB3, IL1R2, ACKR3 |
| 4 | Immune receptor activity | $2.04 \times 10^{-4}$ | F3, FCGR2A, FCGR3B, FPR1, IL18R1, C3AR1, C5AR1, CCRL2, CD44, IL1RL1, OSMR, HLA-DOA, HLA-DQA1, CCR1, LILRA5, LILRA6, IFNGR1, CSF2RA, CSF2RB, IL1R1, IL1RAP, IL4R, IL6R, IL6ST, IL7R, IL15RA, IL1R2, ACKR3 |
| 5 | Extracellular matrix binding | $2.04 \times 10^{-4}$ | SPARCL1, ANXA2, ANXA2P2, SPARC, SPP1, ADAM9, TINAGL1, GPC1, THBS1, ADGRG1, CCN1, CTSS, ITGA3, ITGAV, ITGB1, ITGB3 |
| 6 | Nuclear localization sequence binding | $1.70 \times 10^{-3}$ | RANBP6, NFKBIA, NUP98, NOLC1, KPNA6, NUP153, KPNA4, TNPO1, LBR |
| 7 | Cytokine activity | 0.034 | CCL2, CCL3, CCL4, CCL19, CCL20, AREG, TNFSF11, TNFSF13B, SPP1, BMP2, CXCL2, |
|  |  |  | CXCL3, TNFRSF11B, TGFB3, CLCF1, NRG1, CCL4L1, GDF15, CL4L2, CSF1, IL1B, IL1RN, IL6, CXCL8, INHBA, INHBB, EDN1, LIF, NAMPT |

**Table S7:** Demographics, disease status, and serum biochemistries at the time of collection of PBMC for flow cytometric studies, Related to STAR Methods.

| Patient ID | Gender | Diagnosis | Age | ALT (IU/L) | ALP (IU/L) | GGT (IU/L) | Total Bilirubin (mg/dL) | Direct Bilirubin (mg/dL) |
| --- | --- | --- | --- | --- | --- | --- | --- | --- |
| HIA057 | F | EHBA | 0.63 | 96 | 283 | 63 | 26 | 21.2 |
| HIA061 | M | EHBA | 2.76 | 126 | 908 | 306 | 4.7 | 3.8 |
| HIA065 | M | congenital CMV;<br>EHBA | 0.21 | 169 | 580 | 162 | 18.1 | 14.6 |
| HIA067 | F | EHBA |  | 100 | 314 | 972 | 1.8 | 1.1 |
| HIA063 | M | EHBA | 0.11 | 117 | 330 | 1199 | 5.6 | 5 |
| HIA069 | M | EHBA | 0.59 | 73 | 670 | 298 | 11.6 | 8.2 |
| HIA090 | F | EHBA | 0.13 | 31 | 428 | 1099 | 6.3 | 4.7 |
| HIA092 | F | EHBA | 0.10 | 88 | 583 | 1002 | 7.4 | 5.7 |
| HIA082 | M | healthy control | 0.17 | - | - | - | - | - |
| HIA081 | M | healthy control | 2.89 | 20 | 106 |  | 0.7 | 0.2 |
| HIA091 | M | healthy control | 1.43 | - | - | - | - | - |

**Figure S1.**
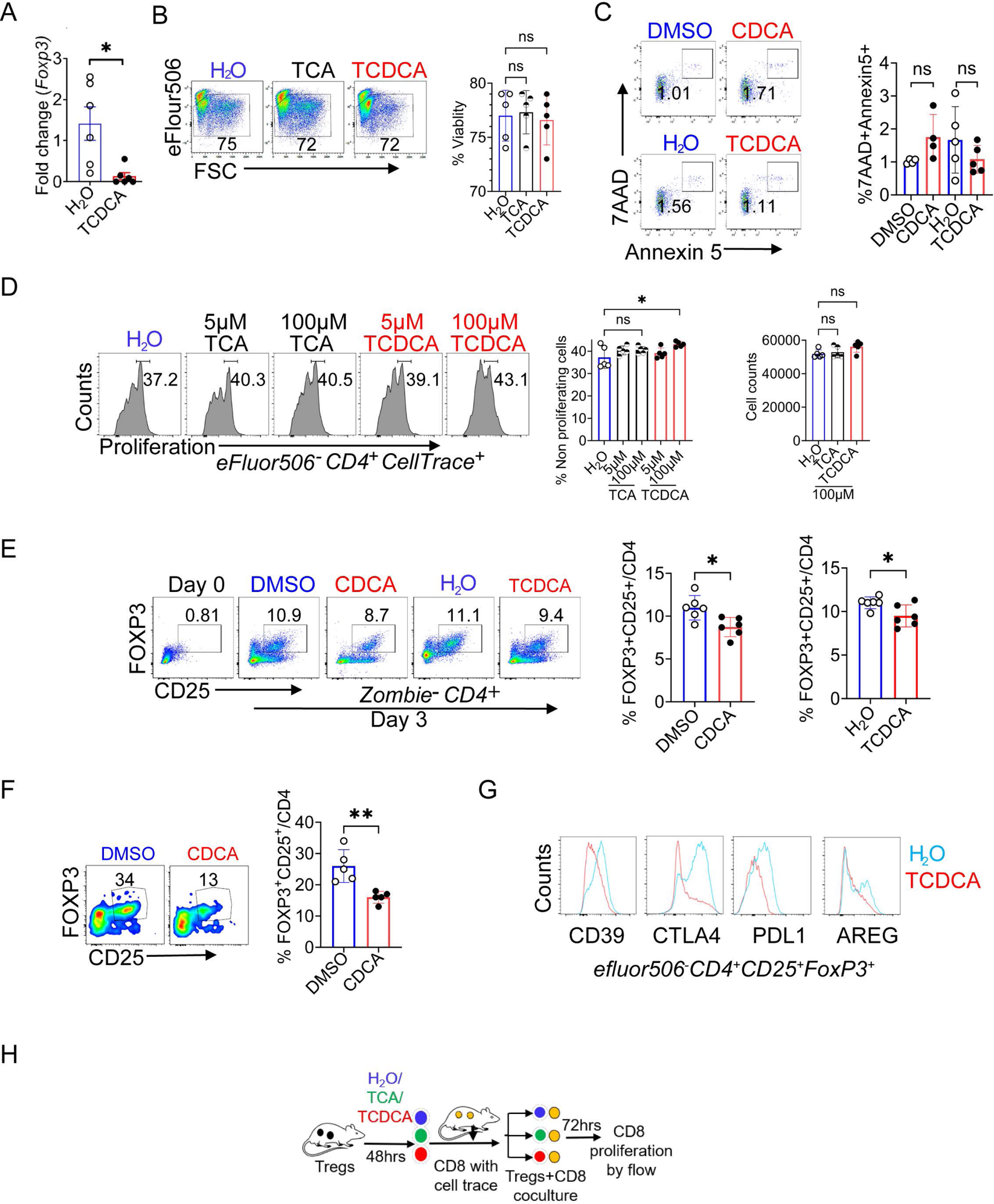
Related to Figure 1. Bile acids repress FOXP3 expression and suppressor function of Tregs. FACS purified splenic GFP+CD25+CD4+ Tregs from FOXP3(EGFP) reporter. Mice were stimulated for 24 hours with 100 µM of TCA, 100 µM of TCDCA, or vehicle (H_2_O), αCD3/CD28 and IL-2 prior to isolation of RNA for RT-qPCR (**A**). Viability of Tregs after 48 hours of stimulation with aCD3/CD28 and culture with IL-2, TCA, TCDCA, or vehicle (H_2_O) was tested with eFluor 506 fixable viability dye for detecting dead cells (**B**). Apoptosis of cultured Tregs following stimulation with 10µM of CDCA, 100µM of TCDCA, or corresponding vehicle was determined by flow cytometry for Annexin 5 and 7AAD (**C**). Proliferative capacity of Tregs stimulated with aCD3/CD28 in media conditioned with IL-2 and TCA or TCDCA in concentrations ranging between 5 and 100µM was assessed CellTrace Violet Cell Proliferation Kit (**D**). Naive splenic CD62L+ CD4 cells were cultured under Treg-polarizing conditions with αCD3/CD28 beads (1:1 ratio), hIL-2 (100U) and TGF-β1 (10 ng/ml), in presence of CDCA (10μM) and TCDCA (100μM) or H_2_O (vehicle) for 72 hours prior to cytometric enumeration of Tregs (**E**). Naïve CD45RA+ T cells from healthy blood donors were cultured with αCD3/CD28 beads (1:1 ratio), hIL-2 (100U) and TGF-β1 (10 ng/ml), in presence of CDCA (10μM) for 72hrs prior to flow cytometry (**F**). Tregs were cultured and stimulated for 48 hours with 100 μM of TCDCA after which expression of the suppressor molecules CD39, CTLA4, PDL1 and AREG was assessed by flow cytometry (**G**). Schematic representation of CD8/Treg co-culture proliferation assay to determine the suppressor function of Tregs pretreated with 100µM TCA, 100 µM TCDCA, or H_2_O (**H**). Unpaired t test except **D** was applied to test for statistical differences between groups, \**p*<0.05 and \*\**p*<0.01.

**Figure S2.**
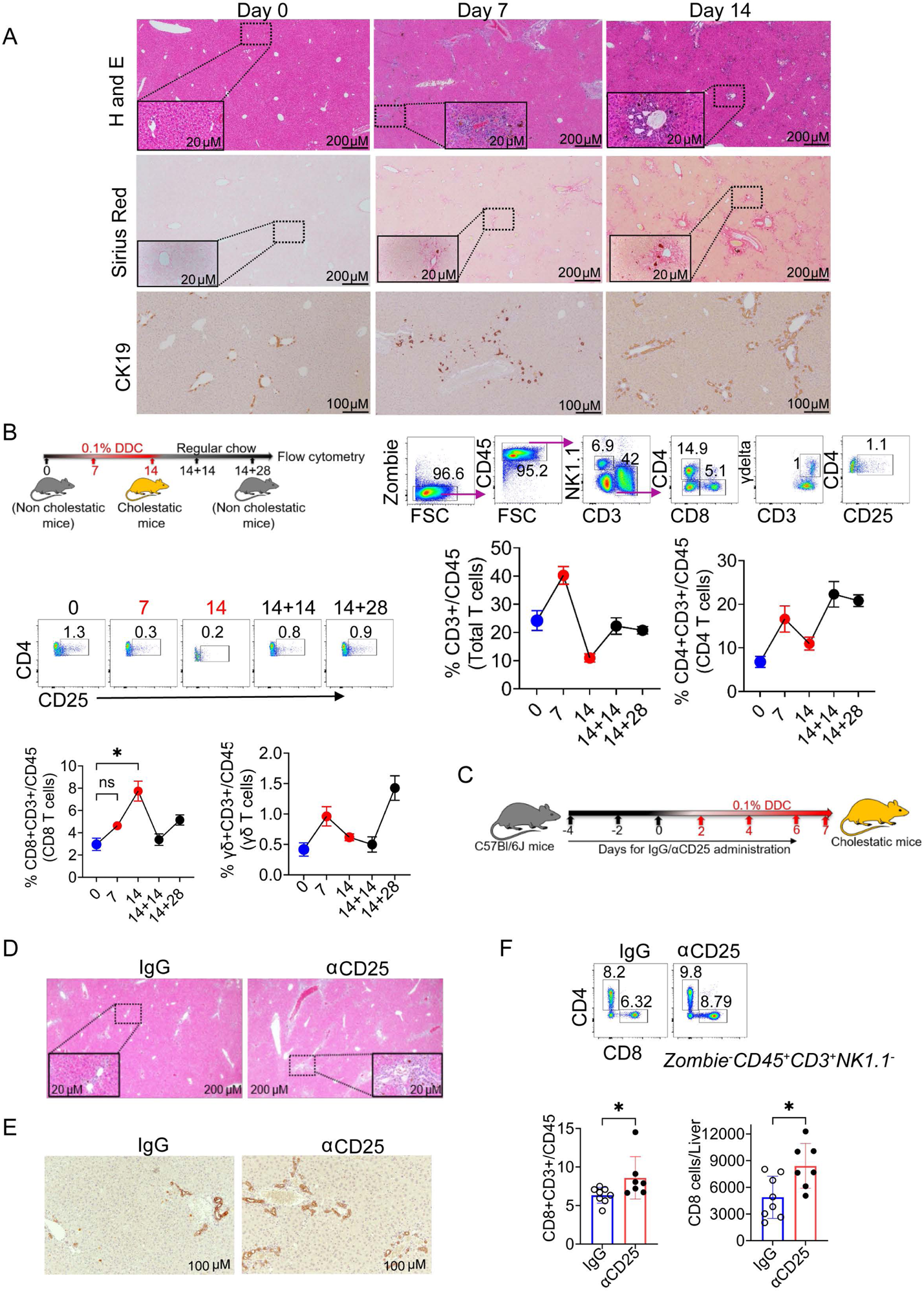
Related to Figure 1. Hepatic immune cell composition during and after DDC-induced cholestatic liver injury. 8-week-old, male C57BL/6 mice were fed with DDC admixed at 0.1 % (w/w) in 5058 Lab chow for 7 or 14 days. Representative photomicrographs from liver sections stained with H&E for inflammatory cell infiltration or Sirius Red for collagen deposition, respectively. Tissue stained for CK19 by immunohistochemistry showed proliferating bile ducts and intrahepatic biliary mass (**A**). Livers from mice at D0, D7, D14, D14+14, and D14+28 of DDC challenge were subjected to multi-parameter flow cytometry to enumerate hepatic lymphocyte composition. Flow plots show the gating strategy for T-cell subsets and percentage of hepatic Tregs in control- and DDC-treated mice (**B**). Experimental design to determine whether CD25+ Tregs protect from DDC induced liver injury. Male C57Bl/6 mice were subjected to αCD25 antibody to deplete Tregs (or isotype matched IgG in controls) prior to challenge with DDC for 7 days (**C**). Representative images from H&E-stained liver sections show increased periportal inflammation in CD25 depleted mice challenged with DDC (**D**). IHC for CK19 highlights increased intrahepatic biliary mass in Treg-depleted mice upon DDC challenge (**E**).

**Figure S3.**
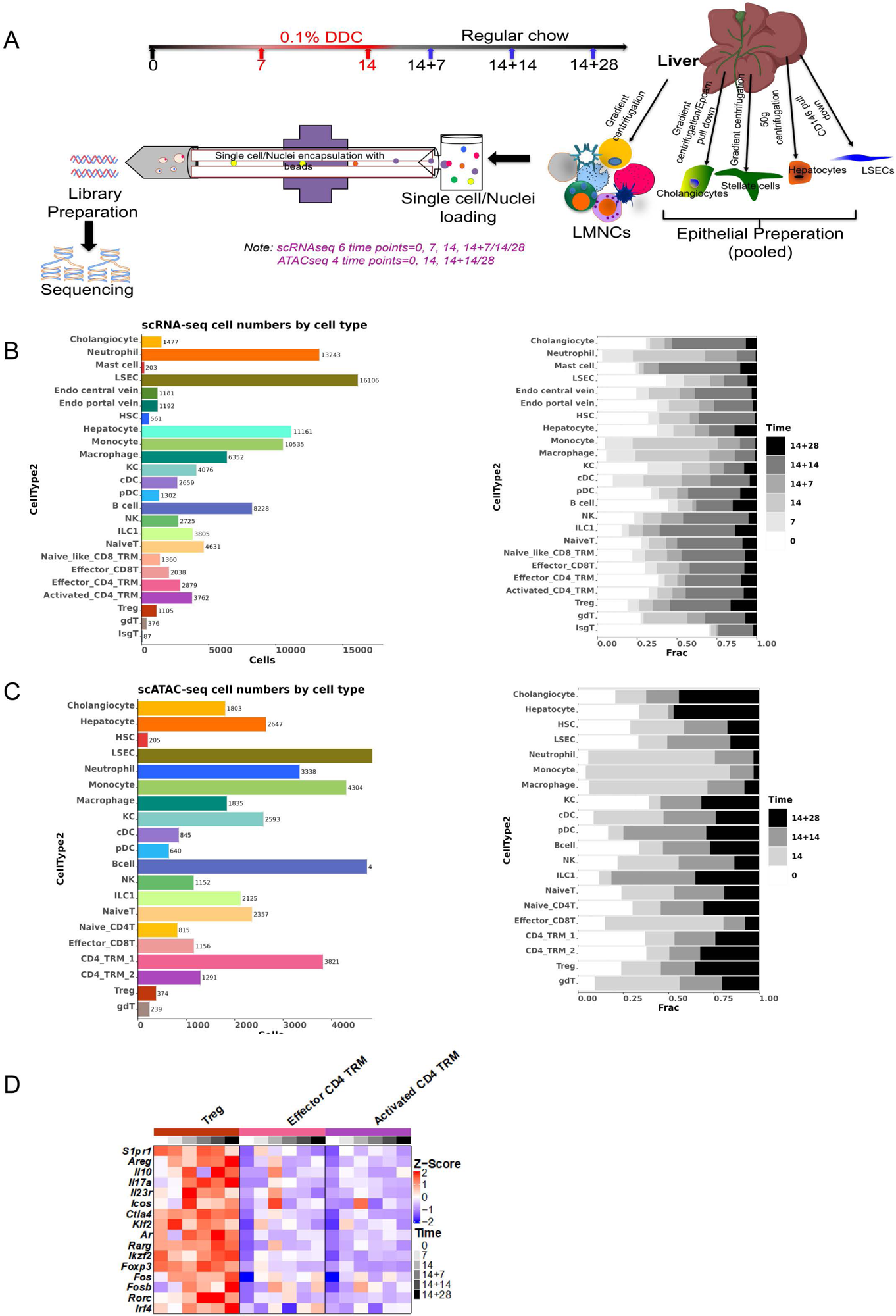
Related to Figure 2. Single-cell genomics of liver epithelial, endothelial, immune cells, stellate cells across time course of DDC-induced sclerosing cholangitis and repair. Schematic representation of the experimental approach for preparation of single-cell suspensions from liver mononuclear, epithelial, endothelial, and hepatic stellate cells and subsequent scRNAseq and snATACseq for mice at baseline (D0), during (D7/14), and following challenge with 0.1% DDC (14+7/14/28) (**A**). Number of single cells passing QC for scRNAseq studies per predicted cell type across six time points of DDC treatment **(B**). Number of single nuclei passing QC for snATACseq studies per predicted cell type across four time points of DDC treatment (**C**). Heatmap of Th17- and Treg-associated genes expressed by Tregs, effector memory T-cells, and activated CD4 lymphocyte populations across the DDC injury course (**D**).

**Figure S4.**
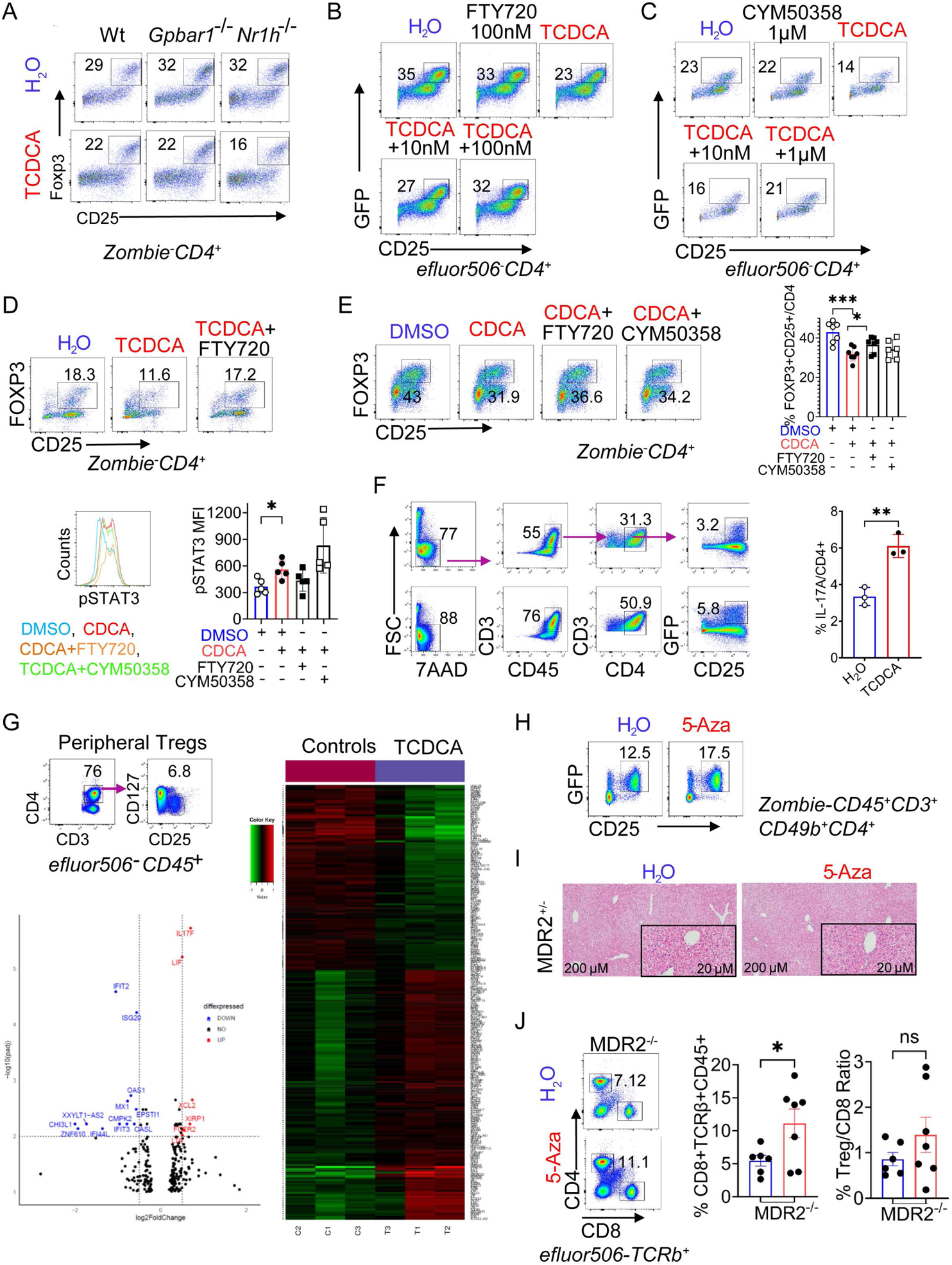
Related to Figure 3. Sphingosine-1-phosphate receptor (S1PR) mediates FOXP3 repression and induction of *Th17* gene expression in Tregs with TCDCA stimulation. Splenic Tregs from wild type (WT), *Gpbar1* (TGR5)^-/-^, and *Nr1h4* (FXR)^-/-^ mice were stimulated with 100μM TCDCA for 48 hrs prior to flow cytometry (**A**). Splenic Tregs from *Foxp3*/GFP reporter mice were cultured and stimulated with 100 μM TCDCA or vehicle along with 10/100nM of the S1PR1 inhibitor FTY720 or with 10nM/1µM of the S1PR4 inhibitor CYM50358 (**B/C/D**). Splenic Tregs stimulated with 10μM CDCA and S1PR1/4 inhibitors for 48 hours prior to cytometric determination of FOXP3 expression or restimulation with PMA/Ionomycin followed by intracellular flow cytometry for pSTAT3 (**E**). Naive splenic CD62L+ CD4 cells from *Il17a*/GFP reporter mice were cultured under Treg-polarizing conditions with αCD3/CD28 (2μg/ml), hIL-2 (100U) and TGF-β1 (10 ng/ml), with TCDCA (100 μM) or H_2_O (vehicle) for 48 hours followed by cytometric enumeration of IL17A (GFP)+ cells (**F**). FACS-sorted human Tregs from blood donors were cultured with 100 µM of TCDCA 48 hrs, after which total RNA was isolated for RNAseq (**G**). Non-cholestatic *Mdr2*^+/-^ mice were treated with 20 µg/g mouse of 5-Aza or vehicle by daily IP injections for seven days prior to cytometric enumeration of hepatic Tregs (**H**). Representative photomicrographs of H&Estained liver sections from livers of *Mdr2*^+/-^ treated with 5-Aza (**I**). Frequency of hepatic CD8 cells was assessed by cytometry in MDR2^-/-^ mice treated with 5-Aza (**J).** Unpaired t-test (**F,** J) or one-way ANOVA with Dunnett multiple comparisons against “CDCA without S1PR inhibitors” in **E** were applied to test for statistical differences between groups with ^ns^non-significant, \**p*<0.05, \*\**p<*0.01, and \*\*\**p*<0.001.

**Figure S5.**
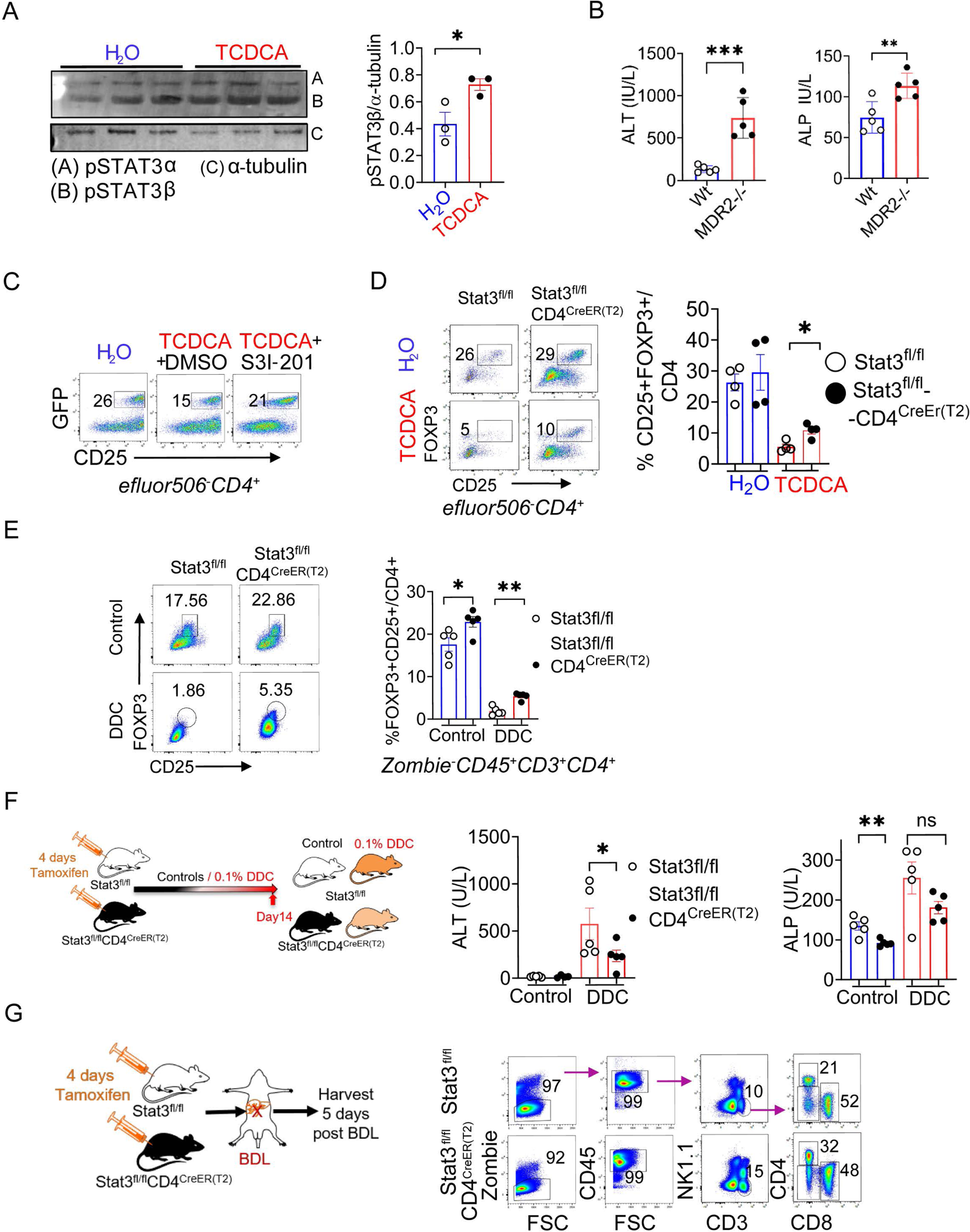
Related to Figure 4. TCDCA-induced repression of FOXP3 in Tregs is associated with induction of STAT3. Splenic Tregs were activated and cultured in presence of TCDCA for 24 hours prior to immunoblotting for phosphorylated α (86 kD) and β isoforms (79 kD)of STAT3 and α-tubulin as loading control followed by densitometry (**A**). Serum samples were collected from 45-day-old WT and *Mdr2*^-/-^ mice and subjected to colorimetric assays for ALT and ALP (**B**). FACS-sorted splenic Tregs from *Foxp3*^EGFP^ reporter mice were cultured with 100µM of TCDCA or H_2_O in media conditioned with 100μM of the STAT3 antagonist S31-201 or DMSO (**C**). *Stat3*^fl/fl^ and *Stat3*^fl/fl^CD4^CreER(T2)^ mice were treated with tamoxifen for 4 days, after which splenic Tregs were isolated and stimulated for 48 hours with 100µM of TCDCA and analyzed by flow cytometry (**D**). MACS sorted splenic Tregs from WT mice were cultured with 100µM of TCDCA or H_2_O in media conditioned with 100µM of S31-201 for 48 hours followed by intracellular flow cytometry for RORγt and IL17A (**E**). *Stat3*^fl/fl^ and *Stat3*^fl/fl^CD4^CreER(T2)^ mice were treated with tamoxifen for 4 days followed by challenge with 0.1% DDC for 14 days prior to collection of serum for ALT and ALP measurement (**F)** or followed by bile duct ligation (BDL) and cytometric enumeration of liver infiltrating inflammatory cells five days later (**G**). Unpaired t-test or one-way ANOVA with Dunnett multiple comparisons against “TCDCA with DMSO” in **E** were applied to test for significant differences between groups with ^ns^non-significant, \**p*<0.05 and \*\**p*<0.01 and \*\*\**p*<0.001.

**Figure S6.**
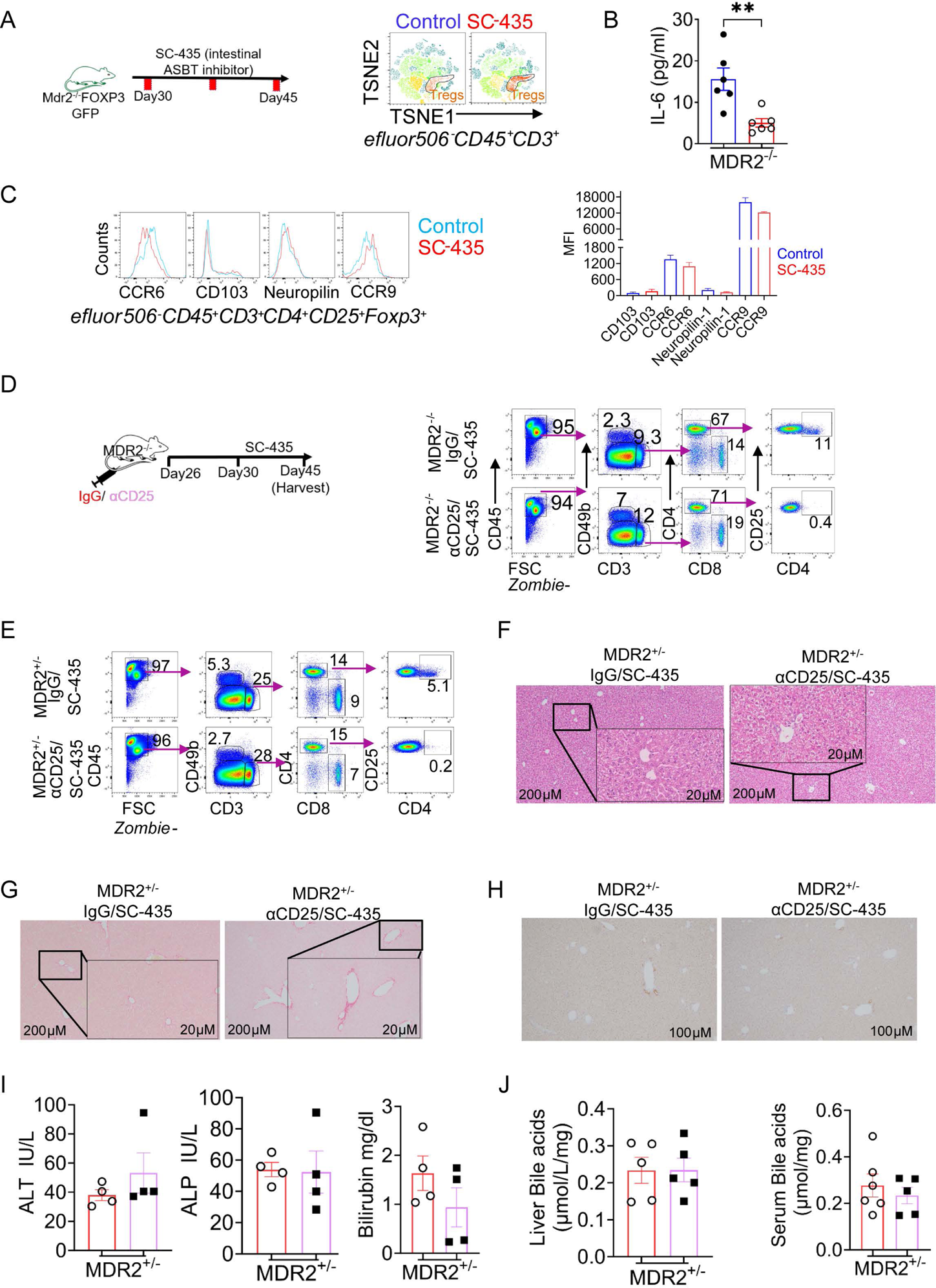
Related to Figure 5. Interruption of intestinal bile acid reabsorption promotes infiltration of the liver with Tregs. 30-day-old female *Mdr2*^-/-^ mice were treated with 0.008% (wt/wt) of the IBATi SC-435 admixed to 5058 Lab diet 14 days prior to isolation and subsequent multiparameter flow cytometry of LMNCs. Data is represented in TSNE plots denoting frequency within CD4 compartments (**A**). End-of-treatment serum samples were subjected to Luminex assay (**B**). *Mdr2*^-/-^ FOXP3 (EGFP) mice were treated with SC-435 (red) or control chow (blue) for 14 days prior to cytometric determination of surface markers associated with gut homing of Tregs (**C**). *Mdr2*^-/-^ mice were simultaneously treated with SC-435 and either αCD25 antibodies (250μg/mice) or IgG control for 2 weeks prior to cytometric enumeration of hepatic Tregs (**D**). 30-day-old, non-cholestatic *Mdr2*^+/-^ mice were simultaneously treated with SC-435 and IgG/aCD25 (250μg/mice) antibodies for 2 weeks prior to cytometric enumeration of hepatic Tregs (**E**), staining of liver sections with H&E or Sirius Red (**F**/**G**), immunohistochemistry for CK19 (**H**), measurement of serum ALT, ALP, and total bilirubin (**I**), and determination of hepatic and serum bile acid concentrations (**J**). Each data point represents an individual mouse. Unpaired t-test was applied to test for significant differences between groups with \**p*<0.05 and \*\**p*<0.01.

**Figure S7.**
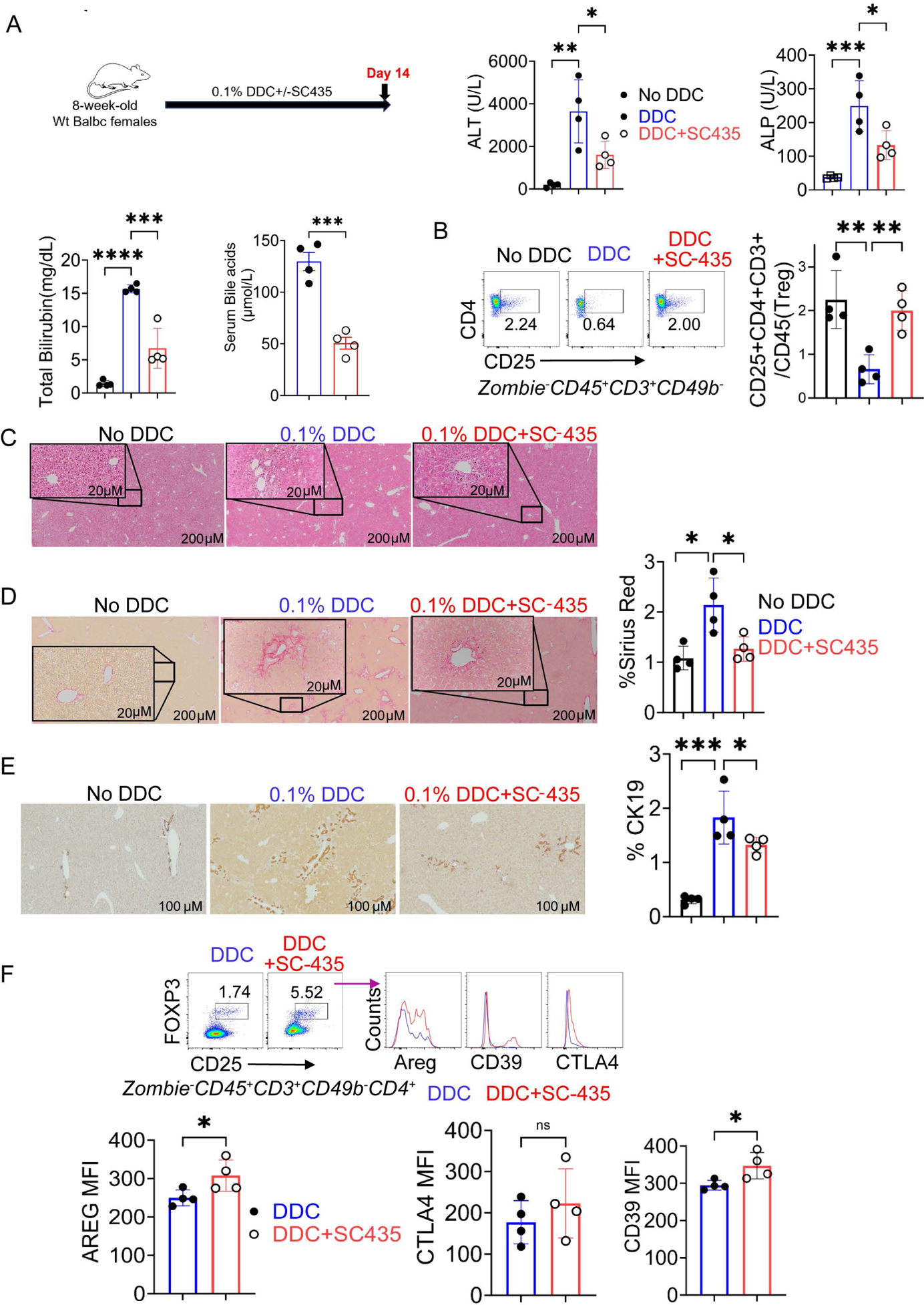
Related to Figure 5. Blockade of intestinal bile acid reabsorption promotes a surge of hepatic Tregs in DDC induced cholestatic liver disease. Wild type mice were fed with either 0.1% DDC to induce SC or DDC chow also containing 0.008% (wt/wt) SC-435 for 14 days prior to measurement of serum biochemistries (ALT, ALP, TB) (**A**), cytometric enumeration of hepatic Tregs and CD8 cells (**B**), assessment of liver histomorphology on liver sections stained with H&E (**C**) and Sirius Red (**D**), and immunohistochemistry for CK19 (**E**). Suppressor molecule expression on hepatic Tregs from mice in the aforementioned treatment groups was assessed by flow cytometry (**F**). One-way ANOVA with Dunnett multiple comparisons against the group “DDC” **in A-E** and unpaired t-test in **F** were applied to test for significant differences between groups with \**p*<0.05 and \*\**p*<0.01 and \*\*\**p*<0.001.

**Figure S8.**
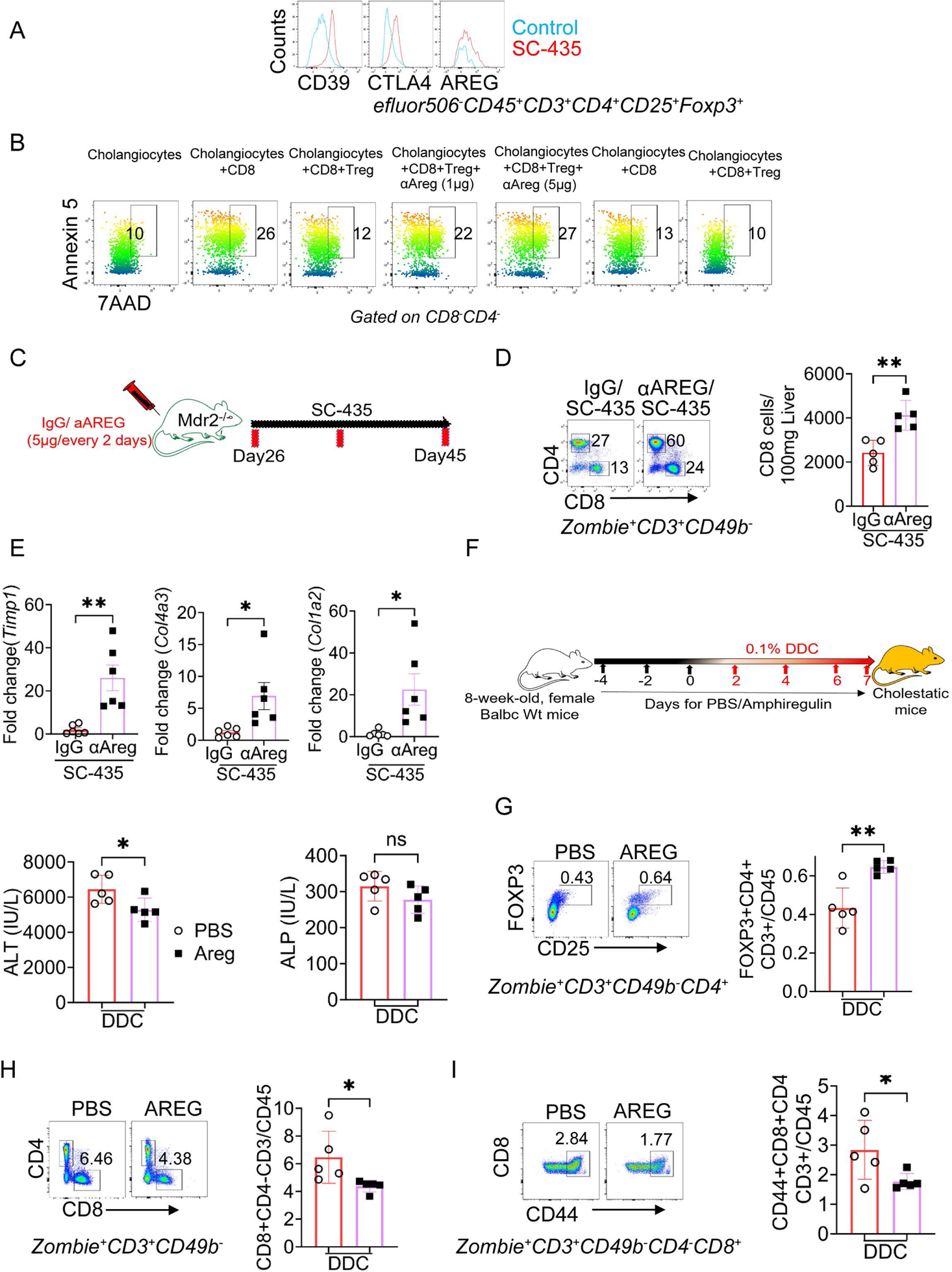
Related to Figure 6. AREG mediates Treg suppression of CD8 cells in sclerosing cholangitis. 30-day-old, female *Mdr2*^-/-^ mice treated with SC-435 (red) or control chow (blue) for 14 days prior to cytometric determination of surface markers associated with suppressor function of Tregs (**A**). Neonatal cholangiocytes were cultured for 24 hours in presence/absence of hepatic Tregs from adult WT mice, CD8 lymphocytes from adult MDR2^-/-^ or WT mice, and 1 or 5 μg of AREG neutralizing antibody. Cholangiocyte death by apoptosis was assessed cytometrically by positive staining for 7AAD and Annexin-5 (**B**). *Mdr2*^-/-^ mice were simultaneously treated with SC-435 and AREG-neutralizing antibody or IgG prior to cytometry enumeration of CD4 and CD8 lymphocytes (**C/D**) and quantitation of hepatic expression for *Timp1*, *Col1a2,* and *Col4a3* by qPCR (**E**). BALB/cJ mice were supplemented with recombinant 5µg/mouse of AREG (PBS control) every two days, starting 4 days before DDC challenge. After treatment course, serum was collected for serum biochemistries (**F**) and flow cytometry was applied to enumerate frequency of Tregs, CD4 and CD8 lymphocytes, and CD44+ effector CD8 cells in LMNCs (**G-I).** Each data point represents an individual mouse. Unpaired t-test was applied to test for significant differences between groups with \**p*<0.05 and \*\**p*<0.01.

**Figure S9.**
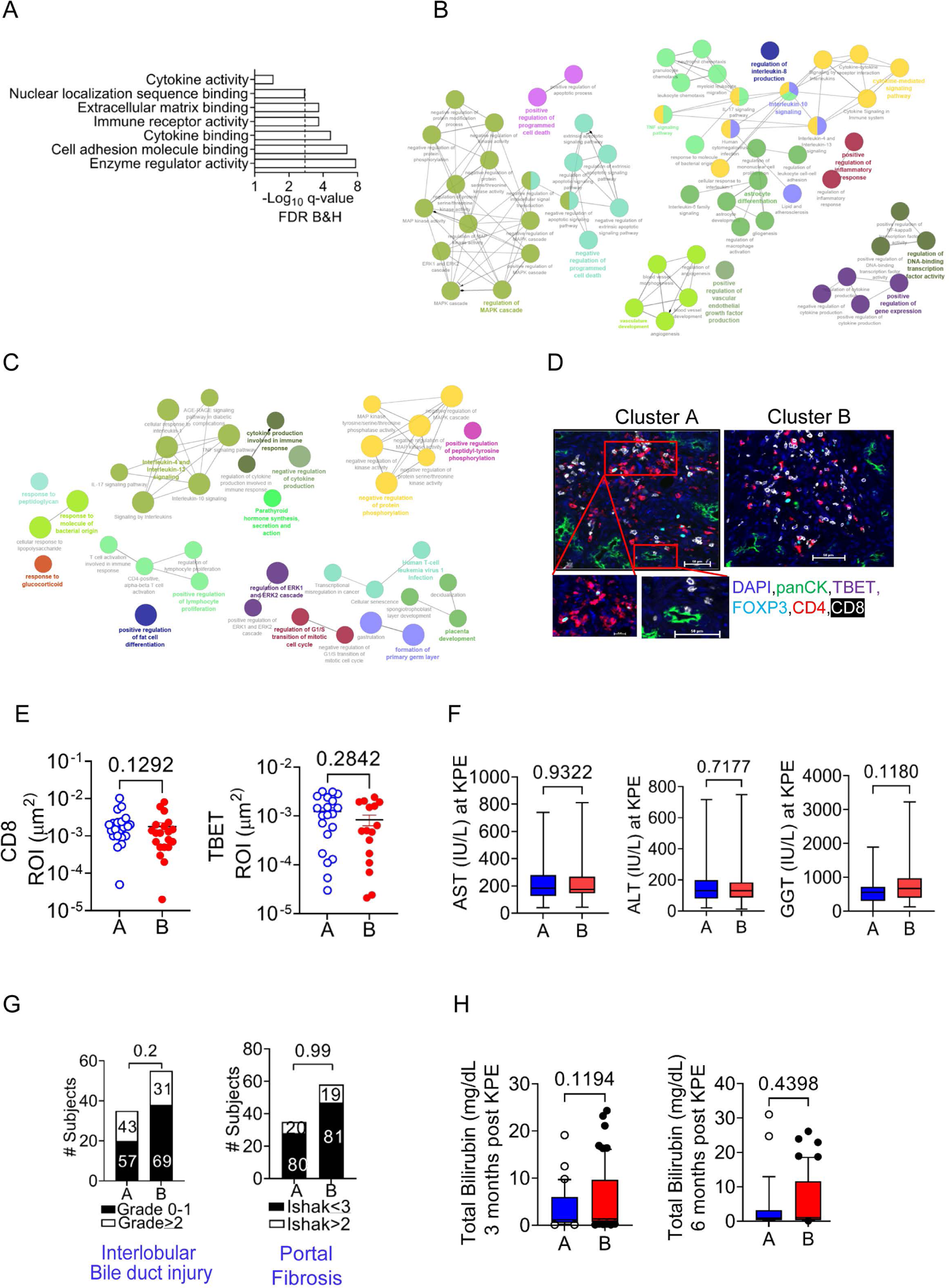
Related to Figure 7. Segregation of patient cluster by Treg-liver transcriptome with clinical phenotype. A set of differentially expressed genes was derived from liver RNAseq data of subjects with “high” vs “low” Treg counts by IF and applied to the entire cohort of infants with liver RNAseq data available from the time of diagnosis with EHBA (n=130) or idiopathic neonatal cholestasis (INH; n=10). Unsupervised analysis identified 2 clusters of patients based on similar expressions of Treg associated genes. The graph displays q-values for GO pathways upregulated in patients in cluster A with increased number of Tregs. Dotted line denotes pathways with significant differences between clusters to the right (**A**). The cytomaps display pathways of upregulated genes in patients with high Tregs (**B)** and gene pathways in patients with upregulated AREG (**C**). Representative images of multi-parameter immunofluorescence DAPI, CD4, FOXP3, panCK, CD8, TBET on sections from liver tissue of diagnostic biopsies in infants with extrahepatic biliary atresia (**D**). Numbers of CD8+ and TBET+ cells per area ROI were determined by automated image analysis of IF in a cohort of 40 patients with EHBA assigned to clusters A and B by liver RNAseq (**E**). Serum AST levels at diagnosis (**F**), grade of interlobular bile duct injury and Ishak stage of portal fibrosis by central review of liver histopathology (**G**), and total bilirubin levels at 3 months and 6-month after KPE in patients assigned to clusters A and B by liver RNAseq (**H**).

